# Reaction mechanisms of Pol IV, RDR2 and DCL3 drive RNA channeling in the siRNA-directed DNA methylation pathway

**DOI:** 10.1101/679795

**Authors:** Jasleen Singh, Vibhor Mishra, Feng Wang, Hsiao-Yun Huang, Craig S. Pikaard

## Abstract

In eukaryotes with multiple small RNA pathways the mechanisms that channel RNAs within specific pathways are unclear. Here, we reveal the reactions that account for channeling in the siRNA biogenesis phase of the Arabidopsis RNA-directed DNA methylation pathway. The process begins with template DNA transcription by NUCLEAR RNA POLYMERASE IV (Pol IV) whose atypical termination mechanism, induced by nontemplate DNA basepairing, channels transcripts to the associated RNA-dependent RNA polymerase, RDR2. RDR2 converts Pol IV transcripts into double-stranded RNAs then typically adds an extra untemplated 3’ terminal nucleotide to the second strands. The dicer endonuclease, DCL3 cuts resulting duplexes to generate 24 and 23nt siRNAs. The 23nt RNAs bear the untemplated terminal nucleotide of the RDR2 strand and are underrepresented among ARGONAUTE4-associated siRNAs. Collectively, our results provide mechanistic insights into Pol IV termination, Pol IV-RDR2 coupling and RNA channeling from template DNA transcription to siRNA guide strand/passenger strand discrimination.

## Introduction

In eukaryotes, RNA-based surveillance systems keep viruses and transposable elements under control in order to minimize the deleterious consequences of genetic invasion, transposition, mutation and chromosome instability (Slotkin and Martienssen, 2007). These surveillance systems can involve multiple small RNA biogenesis pathways generating guide RNAs that, in association with an Argonaute family proteins, basepair with mRNAs to direct their translational inhibition or degradation or associate with chromatin to direct chemical modifications that repress gene transcription (Borges and Martienssen, 2015; Ghildiyal and Zamore, 2009; Holoch and Moazed, 2015).

In plants, multiple RNA silencing pathways defend against invading or selfish nucleic acids, with the dominant process for transcriptional silencing being RNA-directed DNA methylation (RdDM) (Matzke et al., 2015). In the major RdDM pathway (Figure 1A), 24 nt siRNAs interact with longer noncoding RNAs to guide the methylation process (Wendte and Pikaard, 2017). Mutations in numerous genes affect 24 nt siRNA levels, with loss of NUCLEAR RNA POLYMERASE IV (Pol IV), RNA-DEPENDENT RNA POLYMERASE 2 (RDR2) or DICERLIKE 3 (DCL3) being most deleterious (Herr et al., 2005; Onodera et al., 2005; Xie et al., 2004). Once synthesized, 24 nt siRNAs associate with ARGONAUTE4 (AGO 4) (Zilberman et al., 2003), or a related AGO family member, and guide the complexes to target sites transcribed by NUCLEAR RNA POLYMERASE V (Pol V) (Wierzbicki et al., 2008; Wierzbicki et al., 2009). Through basepairing with Pol V transcripts and AGO-Pol V interactions (El-Shami et al., 2007; Wierzbicki et al., 2009), siRNA-AGO silencing complexes mediate recruitment of chromatin modifiers that include the *de novo* DNA methyltransferase, DRM2, which methylates cytosines in the region in all sequence contexts (CG, CHG or CHH, where H is any nucleotide other than G) (Bohmdorfer et al., 2014; Cao and Jacobsen, 2002; Wierzbicki et al., 2009). Histone post-translational modifications occur in cross-talk with DNA methylation, collectively resulting in chromatin environments that suppress promoter-dependent transcription by DNA-dependent RNA Polymerases I, II or III (Du et al., 2015). In the germline of mammals, an analogous pathway involves piRNAs, so-named for their association with proteins of the PIWI subfamily of Argonaute proteins (Ozata et al., 2018), that guide transposon methylation by DNMT3a and DNMT3b, the orthologs of plant DRM2 (Chedin, 2011; Skvortsova et al., 2018).

**Figure 1.**
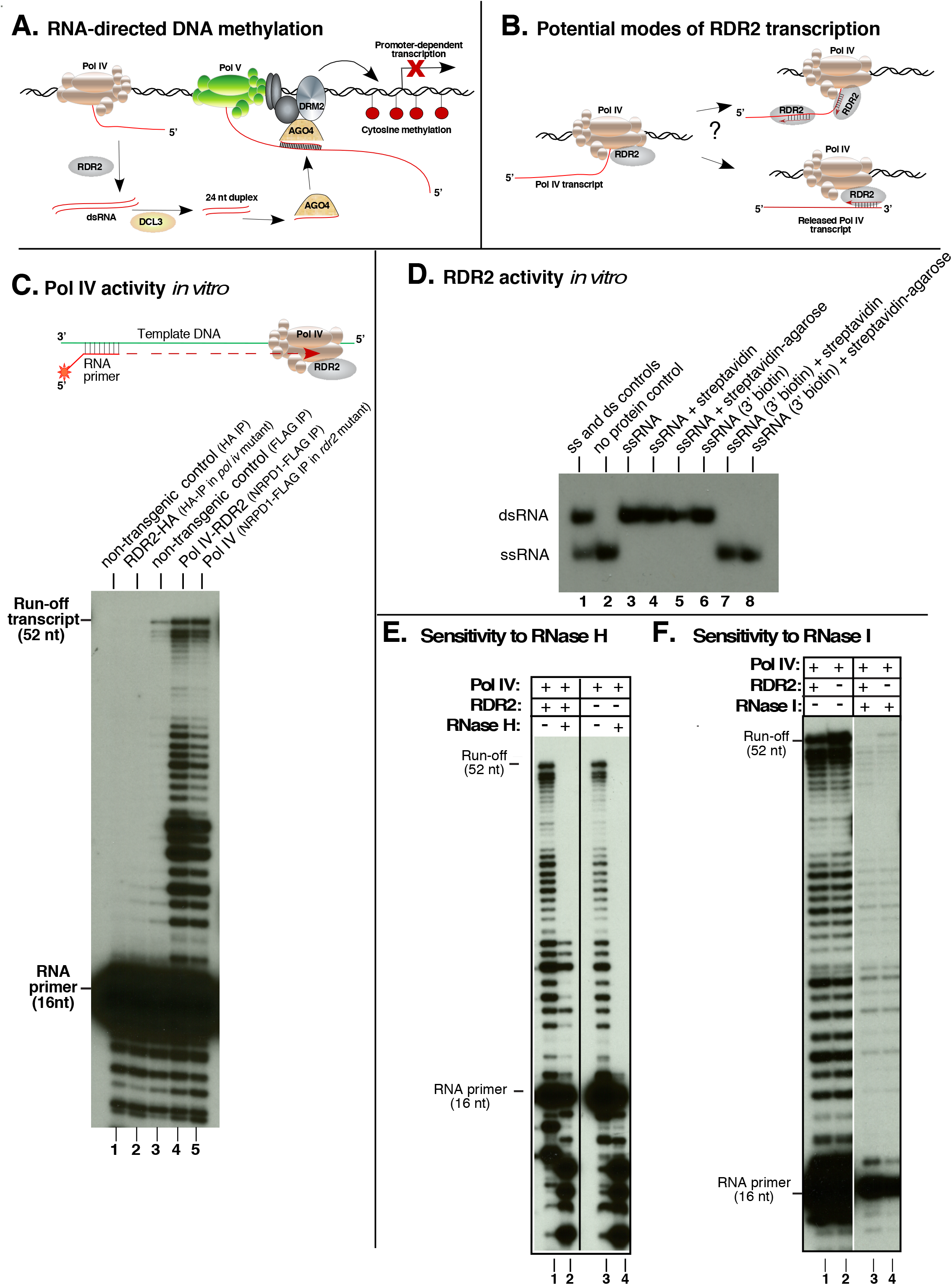
RNA polymerase activities of Pol IV and RDR2. **A.** A simplified model for RNA-directed DNA methylation. **B.** Potential modes of Pol IV-RDR2 cooperation in dsRNA synthesis. **C.** Pol IV transcription is independent of RDR2. Affinity purified Pol IV and/or RDR2 were tested for the ability to extend a 5’ end-labeled 16 nt RNA primer hybridized to DNA oligonucleotide template. Transcripts resolved by denaturing polyacrylamide gel electrophoresis were visualized by autoradiography. HA-tagged RDR2 purified from a Pol IV mutant background *(nrpd1)* was tested in lane 2. FLAG-tagged Pol IV purified with associated RDR2 or from a *rdr2* null mutant was tested in lanes 4 and 5, respectively. Lanes 1 and 3 are controls testing non-transgenic plant lysates subjected to anti-HA or anti-FLAG affinity purification. **D.** RDR2 can convert ssRNA into dsRNA *in vitro*. Purified, recombinant RDR2 was assayed for activity using a 37 nt 5’ end-labeled ssRNA template. Reaction products resolved by nondenaturing PAGE were visualized by autoradiography. Lane 1 shows 37 nt ssRNA and 37 bp dsRNA controls. In lanes 2-5, the ssRNA template tested had a 3’OH group. RDR2 was included in all reactions except for lane 2. Reactions of lanes 4 and 5 included free streptavidin or streptavidin-agarose resin. Reactions of lanes 6-8 tested ssRNAs with a biotin group on the 3’ terminal nucleotide. Streptavidin or streptavidin-agarose were included in lanes 7 and 8. **E.** Pol IV transcripts generated from ssDNA templates are sensitive to RNase H. Pol IV-RDR2 was tested in lanes 1 and 2; Pol IV isolated in the *rdr2* mutant background was tested in lanes 3 and 4. Transcripts in lanes 2 and 4 were subjected to RNase H treatment. Positions of run-off transcripts and the labeled RNA primer are shown. **F.** Pol IV transcripts generated from ssDNA templates are sensitive to RNase I. Pol IV-RDR2 was tested in lanes 1 and 3; Pol IV isolated from *rdr2* mutant plants was tested in lanes 2 and 4. Transcripts in lanes 3 and 4 were subjected to RNase I treatment.

Transcription by multisubunit DNA-dependent RNA Polymerase II (Pol II) is the first step in piRNA or siRNA biogenesis in most eukaryotes (Holoch and Moazed, 2015). However, in plants, 24 nt siRNA biogenesis requires Pol IV (Herr et al., 2005; Onodera et al., 2005), whose 12-subunit composition (Ream et al., 2009) revealed its origin as a specialized form of Pol II. Pol IV acts in close partnership with RDR2, via undefined mechanisms. The two proteins copurify (Haag et al., 2012; Law et al., 2011) and genetic and genomic evidence indicate that both are needed to produce double-stranded (ds) precursors of siRNAs (Blevins et al., 2015; Li et al., 2015; Zhai et al., 2015). These precursors, which average only ~32 bp (Blevins et al., 2015; Zhai et al., 2015), accumulate in *dcl3* mutants and can be cut into 24 nt RNAs *in vitro* upon addition of purified DCL3 (Blevins et al., 2015), indicating that they serve as the immediate precursors of 24 nt siRNAs.

Interestingly, neither strand of siRNA precursor duplexes is detected in *pol IV* or *rdr2* single mutants indicating a codependence whose molecular basis is unknown. Precursor RNA strands tend to begin with a purine (A or G) (Blevins et al., 2015; Zhai et al., 2015), as is common for DNA-dependent RNA polymerases (Basu et al., 2014). However, precursor strands also tend to have pyrimidines at their 3’ ends, such that complementary strands are also expected to have 5’ purines and 3’ pyrimidines. The resulting inability to definitively identify Pol IV or RDR2 transcripts has led us to refer to siRNA-precursors as P4R2 RNAs (Blevins et al., 2015). An intriguing feature of P4R2 RNAs is that their 3’ terminal nucleotides often do not match the corresponding DNA template (Blevins et al., 2015; Wang et al., 2016; Zhai et al., 2015). One hypothesis has suggested that nucleotide misincorporation by Pol IV, especially at methylated cytosines, induces Pol IV termination, explaining both the 3’-terminal DNA-mismatched nucleotides and the short size of Pol IV transcripts (Zhai et al., 2015). However, RDR2 has terminal transferase activity that can add untemplated nucleotides to RNA 3’ ends, suggesting an alternative hypothesis for the mismatched nucleotides (Blevins et al., 2015). Whether RDR2’s terminal transferase activity might act on Pol IV transcripts, RDR2 transcripts, or both is unknown.

Pol IV transcribes single-stranded (ss) DNA but lacks significant activity using sheared double-stranded (ds) DNA *in vitro* (Haag et al., 2012; Onodera et al., 2005). Our current study provides an explanation, showing that when Pol IV is engaged in transcription of a ssDNA strand it terminates within 12-18 nt after encountering dsDNA. Importantly, Pol IV termination induced in this manner is key to channeling the transcript to RDR2, which converts the Pol IV transcript into dsRNA. We show that single-stranded M13 bacteriophage DNA can template siRNA biogenesis *in vitro*, with Pol IV synthesizing first strand transcripts, RDR2 synthesizing the second strands and DCL3 dicing the duplexes into both 24 bp and 23bp siRNAs, as *in vivo*. DNA-mismatched nucleotides are present at precursor and siRNA 3’ ends, as *in vivo*, with sequencing showing these to be hallmarks of RDR2 transcripts, not Pol IV transcripts. Collectively, the reactions of Pol IV, RDR2 and DCL3 are necessary and sufficient for siRNA biogenesis and can account for the short length of P4R2 RNAs, the origin of untemplated 3’ nucleotides, the mechanism of Pol IV-RDR2 coupling and the channeling of RNAs from DNA template transcription to siRNA strand discrimination.

## Results

### Pol IV and RDR2 functions are separable in vitro

Pol IV has long been assumed to transcribe DNA into RNA transcripts that are then made double-stranded by RDR2 (Figure 1A), but definitive evidence is lacking (Haag et al., 2012; Matzke et al., 2015; Pikaard et al., 2012). RDR2 could potentially act co-transcriptionally with Pol IV (Figure 1B, model at upper right). Alternatively, RDR2 might require Pol IV transcripts to be terminated and released, allowing reverse-complementary strand synthesis from end-to-end (Figure 1B, model at lower right).

To investigate the order of Pol IV and RDR2 action, we purified the enzymes from transgenic plants that express FLAG-tagged NRPD1 (the largest subunit of Pol IV) or HA-tagged RDR2 proteins rescuing *nrpd1* or *rdr2* null mutations, respectively (Haag et al., 2012). Pol IV and RDR2 copurify, but affinity purification of Pol IV from a *rdr2* mutant background or RDR2 from a *nrpd1* mutant background allows each enzymes to be isolated free of the other (Haag et al., 2012).

Using a single-stranded DNA template hybridized to an RNA primer, Pol IV will extend the primer in a templated manner when supplied with ribonucleotide triphosphates (Haag et al., 2012). 5’ end-labeling of the primer with ^32^P allows resulting transcripts, resolved by denaturing polyacrylamide gel electrophoresis (PAGE), to be visualized by autoradiography (Figure 1C). Purified RDR2-HA displays no significant DNA-dependent RNA polymerase activity in this assay (Figure 1C, lane 2), resembling the negative controls (lanes 1 and 3). By contrast, Pol IV-RDR2 (purified Pol IV with associated RDR2) generates full-length 52 nt transcripts as well as shorter RNAs (lane 4). Pol IV isolated from a *rdr2* mutant background displays the same activity as Pol IV-RDR2 complexes (lane 5), indicating that Pol IV activity is independent of RDR2.

RDR2 can likewise function independently of Pol IV (Figure 1D). Using recombinant RDR2, expressed from a baculovirus vector in Sf9 insect cells (Blevins et al., 2015), RDR2 transcribes a 37 nt ssRNA template to generate 37bp dsRNA (Figure 1D). To test predictions of the models of Figure 1B, the 3’ end of the ssRNA template was biotinylated and tested as a template in the presence of soluble streptavidin or streptavidin immobilized on agarose beads (Figure 1D). RNAs with unmodified 3’ hydroxyl groups, or 3’ biotin moieties, were both efficiently converted into dsRNA by RDR2 (Figure 1D, lanes 3 and 6). If the RNA was not biotinylated, neither streptavidin nor streptavidin-agarose beads inhibited RDR2 transcription (lanes 4 and 5). However, if the RNA was 3’ biotinylated, both free streptavidin and streptavidin-agarose (lanes 7 and 8) blocked RDR2 activity. These experiments suggest that RDR2 requires RNA templates with free 3’ ends, consistent with the model at the lower right of Figure 1B.

Because Pol IV and RDR2 can function independently, as shown in Figures 1C and D, one might expect double-stranded RNAs to be produced from a single-stranded DNA template when Pol IV and RDR2 are both present. However, ribonuclease sensitivity tests indicate that this is not the case (Figure 1E, F). Ribonuclease H, which specifically degrades the RNA strands of RNA-DNA hybrids, digests full-length transcription products of Pol IV (Figure 1E, lanes 2 and 4), as well as most shorter primer extension products, indicating that most Pol IV transcripts form RNA-DNA hybrids, not dsRNAs. A subset of short Pol IV transcripts that are 21 or 24-26 nt in size are resistant to RNase H when made by Pol IV-RDR2, but not by Pol IV isolated in the *rdr2* mutant background (compare lanes 2 and 4), suggesting that they might be dsRNAs. However, RNase I, which digests ssRNA, but not dsRNA, digests these bands (Figure 1F). We conclude that Pol IV-RDR2 transcription of single-stranded DNA does not produce appreciable levels of dsRNA.

### Pol IV termination is induced by basepaired nontemplate DNA

Pol IV has negligible activity using double-stranded (ds) template DNA (Haag et al., 2012; Onodera et al., 2005), thus we asked what happens when transcriptionally engaged Pol IV encounters dsDNA, using nontemplate DNA strands of varying length to alter the point of encounter (Figure 2). In the absence of nontemplate DNA, transcripts extending to full-length are produced (Figure 2A, lane 1). Upon annealing a 16 nt nontemplate DNA strand, which forms 15 bp with the template (and has a 1 nt 5’ flap) long transcripts are still obtained, but 44-47 nt transcripts, slightly shorter than full-length (52 nt) become more abundant (Figure 2A, lane 3). A nontemplate strand of 28 nt, forming 27 bp with the template, induced abundant transcripts of 36-41 nt (Figure 2A, lane 5) whose 3’ ends are 12-15 nt beyond the point of Pol IV encounter with the nontemplate strand. A 36 nt nontemplate strand, forming a dsDNA region that extends to within 2 nt of the RNA primer, allowed elongation of the primer for ~ 10 nt, but longer transcripts were substantially reduced. Whether full-length transcripts result from nontemplate strand displacement or a failure of template-nontemplate strand annealing is unclear.

**Figure 2.**
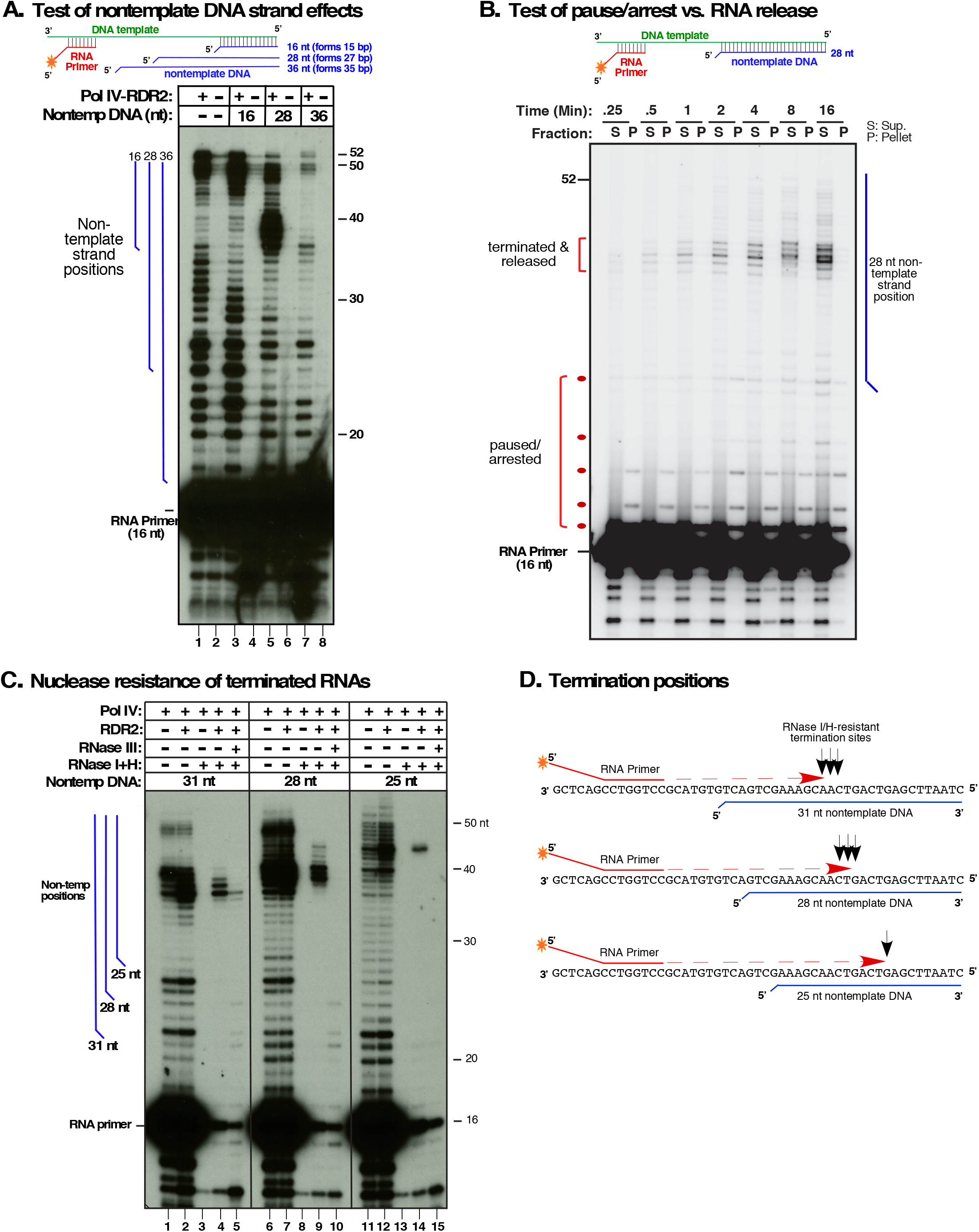
Nontemplate DNA induces Pol IV termination and dsRNA synthesis. **A**. Test of non-template DNA strand effects on Pol IV transcription. Primer-initiated transcription was conducted as in Figure 1C but with nontemplate DNA strands of 16, 28 or 36 nt annealed to the template DNA. Nontemplate DNA strands have an unpaired nucleotide at their 5’ ends. Lanes 1, 3, 5, and 7 show reaction products of Pol IV-RDR2 complexes affinity-purified of FLAG-tagged NRPD1. Lanes 2, 4, 6, and 8 are controls using non-transgenic plant extracts subjected to anti-FLAG affinity purification. Blue vertical lines to the left of the autoradiogram depict the positions of nontemplate strands relative to the transcripts. **B.** Pol IV termination is induced by nontemplate DNA. Primer initiated transcription, using the template −28 nt nontemplate strand pair, was conducted using Pol IV-RDR2 complexes immobilized on anti-FLAG resin. At the indicated timepoints, transcription reactions were subjected to centrifugation and RNAs of supernatant (S) and pellet (P) fractions subjected to denaturing PAGE and autoradiography. **C.** Nontemplate DNA affects termination position and induces production of RNAs with the nuclease sensitivities of dsRNA. Pol IV affinity purified from RDR2 wild-type (+) or *rdr2* mutant (-) backgrounds was tested for primer-initiated transcription of the DNA template annealed to 31nt, 28nt or 25nt nontemplate DNAs. Transcription products were incubated with a mixtures RNAses I and H or with RNase III, as indicated. Blue vertical lines represent the positions of nontemplate strands. **D**. Pol IV termination positions are not sequence-specific. The cartoon shows the positions (black arrows) of the 3’ ends of RNase I/H-resistant Pol IV transcripts relative to the DNA template, RNA primer, and nontemplate DNA strands.

In addition to inducing shortened transcripts, nontemplate DNA strands suppress some transcription products observed in their absence. For instance, RNAs of 32-37 nt observed with the template strand alone are suppressed by the 16 nt nontemplate strand (Figure 2A, compare lanes 1 and 3). The 3’ end of a 37 nt transcript corresponds to the position where the doublestranded DNA region begins, but the 32-36 nt transcripts end prior to where Pol IV encounters the nontemplate strand. Likewise, the 28 nt nontemplate strand suppresses production of transcripts whose 3’ ends map prior to the Pol IV-nontemplate strand encounter point (Figure 2A, compare lanes 1 and 5). These observations suggest that nontemplate DNA strand basepairing may prevent template strand cis-basepairing that affects Pol IV pausing or termination.

To determine if transcripts induced by nontemplate DNA result from Pol IV termination, pausing, or arrest, we tested whether transcripts are released from, or remain associated with, Pol IV. Using Pol IV-RDR2 complexes immobilized on anti-FLAG resin (by virtue of FLAG-tagged NRPD1), transcription was conducted using end-labeled primer RNA and the DNA template annealed to the 28 nt nontemplate strand. At varying times, reactions were subjected to brief centrifugation and resin-associated (pellet, P) and supernatant (S) fractions were subjected to denaturing PAGE and autoradiography (Figure 2B). Nontemplate strand-induced 36-41 RNAs were detected within 15-30 seconds, accumulated throughout the time-course, and were found exclusively in supernatant fractions, indicative of terminated, released transcripts. By contrast, shorter RNAs detected in the pellets are interpreted to be paused or arrested transcripts still associated with Pol IV.

To further test how Pol IV termination sites relate to dsRNA encounter sites, we conducted transcription assays using the template strand annealed to 31, 28 or 25 nt nontemplate DNAs (Figure 2C). The 31 nt nontemplate strand, extending nearest to the RNA primer, induced terminated Pol IV transcripts of 35-38 nt (lanes 1 and 2), the 28 nt nontemplate strand induced 36-41 nt transcripts (lanes 6 and 7) and the 25 nt nontemplate strand induced 40-44 nt transcripts (lanes 11 and 12). In each case, termination occurs 12-16 nt beyond the point where Pol IV first encounters the nontemplate strand, at sites showing no obvious sequence similarity (Figure 2D).

Nuclease sensitivity tests provided the first indication that nontemplate DNA induces Pol IV-RDR2 complexes to synthesize dsRNA. In the absence of RDR2, Pol IV transcripts induced by the 25, 28 or 31nt nontemplate strands are sensitive to a mixture of RNase I and RNase H (Figure 2C, lanes 3,8,13), consistent with the transcripts being ssRNAs or ssRNA-DNA thybrids. However, Pol IV transcripts generated in the presence of RDR2 are resistant to the nucleases, consistent with being strands of dsRNA (lanes 4, 9, 14). Moreover, Pol IV-RDR2 transcripts resistant to RNases I and H are digested by *E. coli* RNase III, which specifically degrades dsRNA (lanes, 5, 10 and 15).

### Detection of second strand synthesis by direct labeling of RDR2 transcripts

In the assays of Figures 1 and 2, the first RNA strand, synthesized by Pol IV, has the labeled 5’ monophosphate of the primer. If a complementary strand is synthesized by RDR2, it should have a 5’ triphosphate, making it a substrate for GTP addition by capping enzyme. To test this hypothesis, RNAs labeled by virtue of the ^32^P end-labeled primer (Figure 3A, lanes 1-8) were compared to RNAs initiated using unlabeled primer but then incubated with capping enzyme and α-^32^P-GTP (Figure 3A, lanes 9-12). As in Figure 2, labeled primer extension products are synthesized by Pol IV, independent of RDR2 (Fig. 3, compare lanes 1 and 2), and terminate early in the presence of the 28 nt nontemplate DNA strand (lanes 3 and 4). Only if RDR2 is present are RNAs induced by the nontemplate strand resistant to RNAses I and H (Figure 3A, compare lanes 7 and 8), indicative of dsRNA. Moreover, RDR2-dependent transcripts can be labeled by ^32^P-GTP capping (lane 12) and are resistant to Terminator Exonuclease (Lucigen Corporation), which degrades RNAs with 5’ monophosphate but not triphosphate groups (<Figure S1).

**Figure 3.**
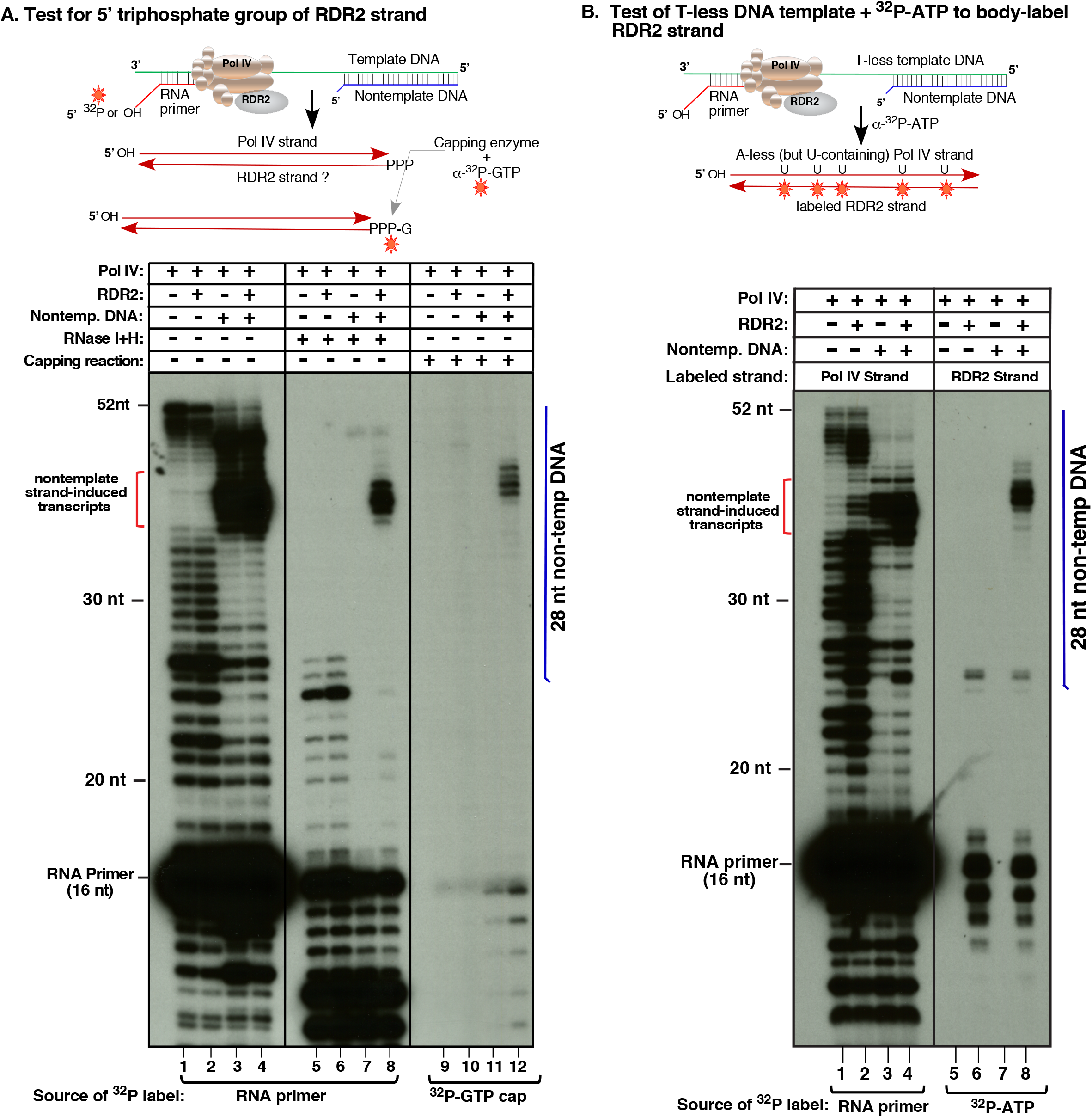
Tests of second strand synthesis by RDR2. **A.** Capping assay to test for a 5’ triphosphate on RDR2-dependent transcripts. First strands, synthesized by Pol IV associated with RDR2 (+) or purified from a *rdr2* mutant (-), were initiated using an RNA primer with a labeled 5’ monophosphate (lanes 1-8) or 5’ hydroxyl group (lanes 912). In half of the reactions, the template was annealed to the 28 nt nontemplate DNA strand to induce Pol IV termination; remaining reactions lacked the nontemplate strand. Lanes 1-4 differ from lanes 5-8 with respect to RNase I/H treatment. In lanes 9-12, vaccinia virus capping enzyme catalyzed labeling with α-^32^P-GTP. **B.** Use of a T-less DNA template and α-^32^P-ATP incorporation to label second strands. All reactions contain Pol IV, associated with RDR2 (+) or purified from an *rdr2* mutant (-) background. In the reactions of lanes 3, 4, 7 and 8, the T-less template was annealed to a 28nt nontemplate DNA strand whose relative position is depicted by the vertical blue line. Transcripts in lanes 1-4 were initiated using a 5’ end-labeled RNA primer to label Pol IV transcripts. In lanes 5-8, transcripts were labeled by α-^32^P-ATP incorporation.

As an independent test of second-strand synthesis by RDR2, a DNA template that lacks thymidines was used to generate transcripts in the presence of α-^32^P-ATP (Figure 3B). With no T’s in the template, A’s are not incorporated into the initial Pol IV strand. However, U’s in the first strand template A incorporation into the second strand (see Figure 3B diagram). Using the T-less template alone, Pol IV generates full length as well as shorter RNAs (lanes 1 and 2). Annealing of a 28 nt nontemplate strand induces early Pol IV termination, generating prominent 37-40 nt transcripts (lanes 3 and 4). If RDR2 is present, the nontemplate-induced 37-40 nt transcripts are converted into dsRNAs, such that ^32^P-ATP is incorporated into the RDR2 strands (compare lanes 5-8). The slower mobility of RDR2 strands, compared to Pol IV strands is explained, in part, by their different sequences and molecular masses (see Figure S2).

Collectively, the results of Figure 3 indicate that Pol IV termination induced by a nontemplate DNA strand is coupled to second strand RNA synthesis by RDR2.

### The extent of template-nontemplate basepairing is critical for Pol IV termination-RDR2 coupling

Multiple variables could potentially affect Pol IV-RDR2 coupling. To test the importance of Pol IV transcript length, we initiated transcription with RNA primers that had 5’ tails ranging from 8 nt to 20 nt (Figure 4A). Each primer yielded transcripts that terminated at the same template positions, generating dsRNAs resistant to RNAses I and H (lanes 2,4,6,8), indicative of Pol IV-RDR2 coupling. These results indicate that RNA transcript length does not dictate positions of Pol IV termination.

**Figure 4.**
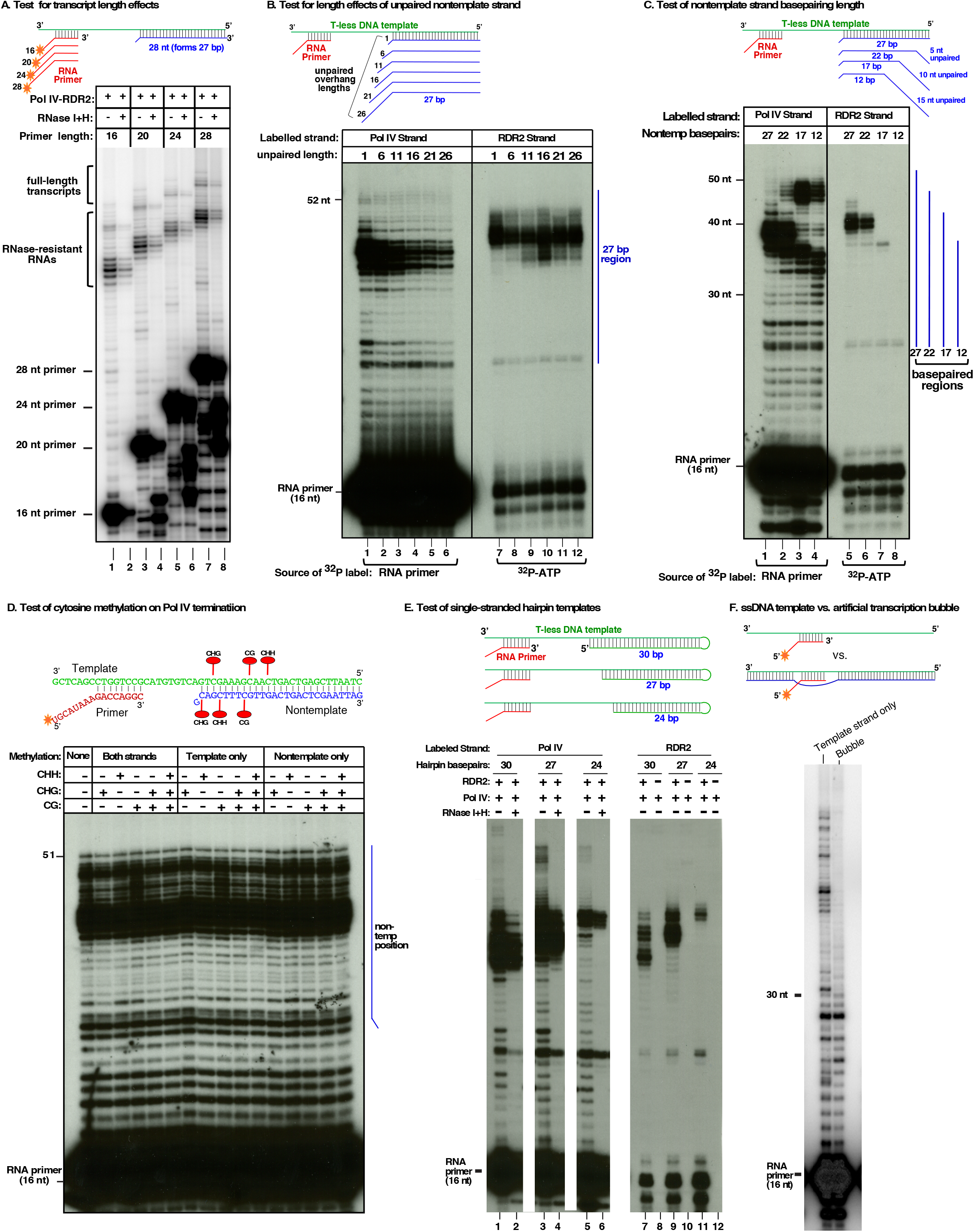
Tests of RNA and DNA parameters for effects on Pol IV termination and RDR2 coupling. **A**. Test of transcript length on Pol IV termination/RDR2 coupling sites. End-labelled RNA primers of 16-28 nt, varying at their 5’ ends (see diagram), were used to initiate transcription by Pol IV-RDR2. One of each duplicate reaction was subjected to RNase I/H digestion. **B**. Test of displaced nontemplate DNA length on Pol IV termination/RDR2 coupling site. Nontemplate DNA strands of 28-53 nt, with variable unpaired 5’ end lengths (1-26 nt), were annealed to the T-less template (see diagram) and tested in Pol IV-RDR2 transcription reactions initiated using an RNA primer. In the reactions of lanes 1-6, the primer was 5’ end-labelled, allowing Pol IV transcript detection. In the reactions of lanes 7-12 the primer was unlabeled and α-^32^P-ATP was used to body-label second strand transcripts. The vertical blue line denotes the 27 basepaired region common to all template-nontemplate combinations. **C.** Test of the number of basepaired nontemplate nucleotides distal to the Pol IV encounter position. 28 nt nontemplate DNA strands forming 12-27 bp with the T-less template, but varying at their 3 ‘ ends, were tested in Pol IV-RDR2 transcription reactions initiated using an RNA primer. In the reactions of lanes 1-4, the primer was 5’ end-labelled, allowing Pol IV transcripts to be visualized. In the reactions of lanes 5-8 the primer was unlabelled and α-^32^P-ATP was used to body-label second-strand RDR2 transcripts. **D**. Test for effects of template and/or nontemplate DNA cytosine methylation on Pol IV termination. Template and nontemplate strands that were unmethylated (far left lane), or methylated at CG, CHG or CHH (in 5’ to 3’ orientation) positions indicated in the cartoon at top, were tested in Pol IV-RDR2 transcription reactions initiated using a 5’ end-labelled primer to visualize Pol IV transcripts. **E.** Test of single-stranded hairpin templates for Pol IV termination/RDR2 coupling. The T-less template was lengthened to allow formation of hairpins with 24-30 bp stems, as indicated in the cartoon. Pol IV-RDR2 transcription reactions were initiated using a 5’ end-labeled primer to visualize Pol IV transcripts (lanes 1-6) or using an unlabeled primer and α-^32^P-ATP to body-label RDR2 transcripts (lanes 7-12). Pol IV purified from *rdr2* null mutant plants was tested in lanes 8, 10 and 12. Reaction products of lanes 2, 4 and 6 were treated with RNases I and H. Images shown are lanes from the same gel and autoradiogram exposure. **F.** Test for Pol IV transcription from an initiation bubble. Primer-initiated Pol IV-RDR2 transcription reactions were conducted using single-stranded template DNA or template DNA annealed to a partially non-complementary DNA strand, as depicted in the cartoon. The primer was 5’ end-labelled to visualize Pol IV transcripts.

We next asked whether the length of displaced DNA of the non-template strand affects Pol IV termination and/or RDR2 coupling. The T-less template was used (as in Fig. 3B), annealed to a set of nontemplate strands that each form 27 bp with the template but have unpaired 5’ ends ranging from 1-26 nt in length (Figure 4B). End-labeled RNA primer was used to detect Pol IV strands (lanes 1-6) snd α-^32^P-ATP incorporation was used to label to label second strands made by RDR2 (lanes 7-12). No effect was observed on the positions of Pol IV termination nor on Pol IV-RDR2 coupling.

To test whether the length of the double-stranded region matters, we tested 28 nt nontemplate strands that form 27, 22, 17 or 12 bp with the T-less template (Figure 4C). Nontemplate strands forming 27 or 22 bp strongly induced Pol IV termination, generating 35-40 nt transcripts terminated 12-17 nt prior to the end of the template (lanes 1 and 2). Reducing basepairing to 17 or 12 bp resulted in longer readthrough transcripts of 46-50 nt, with 35-40 nt transcripts greatly reduced in abundance (lanes 3 and 4). The transcript profile for the reaction using the nontemplate strand that forms only 12 bp with the template resembles the pattern obtained using template DNA in the absence of nontemplate DNA (compare lane 4 to Figure 3B, lane 2).

Using α-^32^P-ATP to label second strands synthesized by RDR2 revealed that abundant RDR2 transcripts are only detected for template-nontemplate pairs that formed 27 or 22 bp (Figure 4C, lanes 5 and 6). Collectively, the results of Figure 4C indicate that the doublestranded region needs to be longer than 17 bp to induce Pol IV termination and Pol IV-RDR2 coupling.

To test whether DNA cytosine methylation affects Pol IV termination, we tested template and/or nontemplate strands that were unmethylated, methylated at individual CG, CHG or CHH motifs, or methylated at all test site motifs (Figure 4D; see diagram for methylcytosine positions). Cytosine methylation had no effect on Pol IV transcription or termination (Figure 4D).

We next asked whether Pol IV termination, and its coupling to RDR2 transcription, requires separable template and nontemplate strands of DNA, or merely basepaired DNA. For this test, we lengthened the T-less template to allow stem-loop formation, generating basepaired stems of 30, 27 or 24 bp, thus recapitulating the experiment of Figure 2C using a single DNA molecule (Figure 4E). The results essentially mirror those of Figure 2C, with the length of Pol IV transcripts determined by the point of encounter with dsDNA.

In the context of double-stranded chromosomal DNA, Pol IV presumably initiates within a locally melted region of duplex DNA, as is true for other multisubunit RNA polymerases, from bacteria to eukaryotic Pols I, II, and III (Bae et al., 2015; Barnes et al., 2015; Holstege et al., 1997; Kahl et al., 2000). To test Pol IV’s ability to carry out transcription in the context of a DNA bubble (Figure 4F), we annealed template and non-template strands, each with 12 nt of non-complementarity, and hybridized an RNA primer to one strand within the resulting bubble, leaving 4 nt between the 3’ end of the primer and the beginning of the double-stranded DNA region. Using the bubble template, Pol IV extends the primer only 14–16 nt whereas in the absence of the nontemplate strand, long transcripts are produced (Figure 4F). These results are consistent with prior results showing that Pol IV extends only ~12-18 nt beyond the point at which it encounters dsDNA.

### Pol IV, RDR2 and DCL3 are sufficient to reconstitute 24 nt siRNA biogenesis in vitro

Using recombinant FLAG-tagged DCL3 produced in insect cells (Figure 5A), we tested whether Pol IV-RDR2 transcripts induced by the 28 nt nontemplate DNA strand are substrates for DCL3 (Figure 5B). Pol IV-RDR2 transcripts, whose Pol IV strands are 5’ end-labeled, are resistant to RNases I/ H (Figure 5B, lane 2) but degraded by RNase III (lane 3), indicative of double-strandedness. Addition of DCL3 converted the Pol IV-RDR2 transcripts into 24 nt RNA products (lanes 5-9).

**Figure 5.**
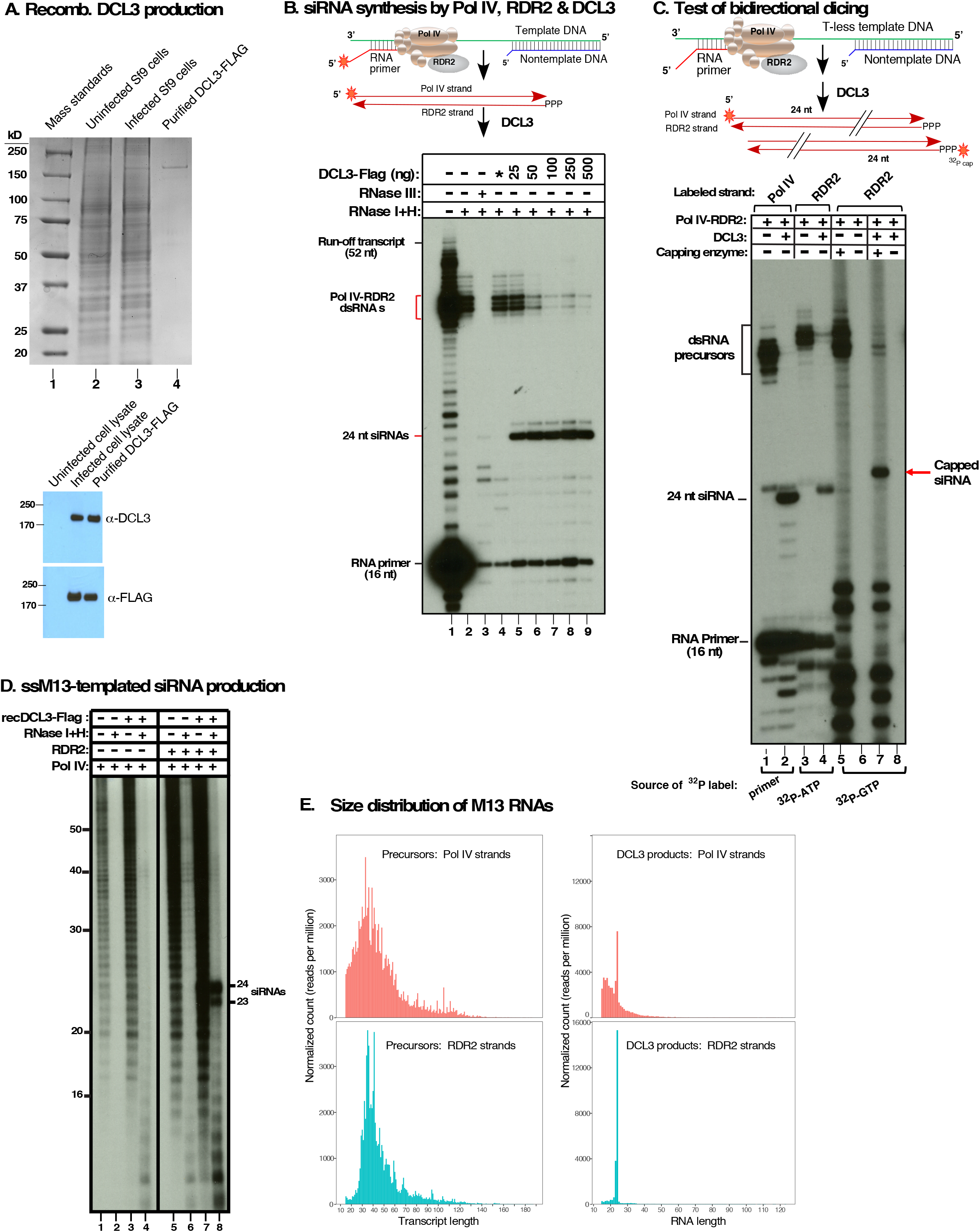
Pol IV, RDR2 and DCL3 are sufficient for siRNA biosynthesis *in vitro*. **A.** Baculovirus-mediated production of recombinant DCL3. Top panel: Stained SDS-PAGE gel showing protein standards (lane 1) adjacent uninfected Sf9 cells lysate (lane 2), DCL3-baculovirus infected cells (lane 3) or affinity purified DCL3-FLAG. Bottom panel: immunoblots using anti-DCL3 or anti-FLAG antibodies. **B.** siRNA synthesis by Pol IV, RDR2 and DCL3. Template DNA was annealed to a 28 bp nontemplate strand and Pol IV-RDR2 transcription reactions initiated from an end-labeled RNA primer. The autoradiogram shows untreated (lane 1), RNase I/H (lane 2), RNase III (lane 3) or DCL3-FLAG digested (25-500 ng; lanes 5-9) transcription products. Uninfected Sf9 cell lysate subjected to anti-FLAG affinity purification serves as a control in lane 4. **C.** Pol IV-RDR2 dsRNA products are diced from either end. Using the T-less template, primer-initiated Pol IV-RDR2 reactions were labeled using end-labeled primer to detect Pol IV strands, using α-^32^P-ATP incorporation to body-label RDR2 strands, or using capping enzyme and α-^32^P-GTP to label RDR2 strand 5’ ends. For each labeling scheme, total transcription products and DCL-diced products are compared. **D.** Pol IV, RDR2 and DCL3 produce siRNAs from single-stranded M13 bacteriophage DNA. Transcription reactions using α-^32^P-ATP to body-label transcripts were conducted using Pol IV-RDR2 (lanes 5-8) or Pol IV purified from *rdr2* mutant plants (lanes 1-4). Transcripts in even lanes were subjected to digestion with RNases I and H. Transcripts in lanes 3, 4, 7 and 8 were incubated with DCL3. **E.** Size distribution and frequency of Pol IV (pink) and RDR2 (blue) strands of precursors and DCL3-diced siRNAs synthesized from M13 template DNA.

To examine the directionality of DCL3 dicing, we used the T-less template to generate dsRNAs that were 5’ end-labeled on the Pol IV strand, body-labeled throughout the RDR2 strand, or 5’ end-labeled on the RDR2 strand, after dicing, by ^32^P-GTP capping (Figure 5C). Labeled 24 nt DCL3 products were observed in each case, showing that DCL3 can process precursor duplexes from either end.

### In vitro biosynthesis of siRNAs from single-stranded M13 DNA

Pol IV transcription initiated with an RNA primer is useful for generating transcripts with defined 5’ ends, but Pol IV does not require a primer and will initiate *de novo* on single-stranded DNA (Haag et al., 2012), including bacteriophage M13 (+) strands (Blevins et al., 2015). This prompted us to test whether Pol IV, RDR2 and DCL3 are sufficient to generate siRNAs from M13mp18 (+) DNA (Figure 5D), whose ~7.3 kb circular genome is predicted to form extensive secondary and tertiary structures that might facilitate Pol IV-RDR2 coupling (Figure S3). Pol IV transcription of M13 DNA, monitored by α-^32^P-ATP incorporation, yields a ladder of transcripts (Figure 5D, lane 1) that are sensitive to RNAse I/ H (lane 2) and are not substrates for DCL3 (lanes 3 and 4). However, if RDR2 is present, a portion of the labeled Pol IV transcripts becomes resistant to RNases I and H (compare lanes 5 and 6), and can be diced by DCL3 into 24 and 23 nt products (lane 8) in a ratio resembling their relative abundance *in vivo* (Kasschau et al., 2007). The molecular basis for producing 24 versus 23 nt siRNA strands is currently unknown.

Because M13 is single-stranded, yet is transcribed into dsRNAs by Pol IV and RDR2, the polymerases responsible for sense and antisense transcripts, relative to DNA template polarity, can be easily ascertained. Deep sequencing of RNAs made by Pol IV alone (isolated from a *rdr2* mutant) and RNAse I/H-resistant RNAs made by Pol IV-RDR2, before and after DCL3 dicing, shows that Pol IV makes first-strand transcripts whose polarity is opposite that of the DNA template (Figures 5E, S4). Only if RDR2 is present are second-strand transcripts produced. These RDR2-dependent transcripts match the polarity of the ssDNA template and thus cannot be DNA-encoded; instead, they can only be generated by transcription of first-strand RNAs made by Pol IV. More than 75,000 unique RNAse I/H-resistant RNAs generated by

Pol IV-RDR2 were sequenced. The size distribution of Pol IV and RDR2 strands is similar, with most being 30-50 nt in length (Figure 5F, left plots). However, more transcripts shorter than 30 nt were detect for Pol IV than for RDR2, suggesting that RDR2 coupling is inefficient if Pol IV transcripts are shorter than ~30 nt.

Sequencing of DCL3-diced products revealed a major peak at 24 nt and a shoulder at 23 nt for both Pol IV and RDR2 strands (Figure 5E, right plots) in keeping with the gel image of Figure 5D (lane 8). RNAs shorter 23 nt are presumably transcripts too short to be diced.

### Untemplated 3’ terminal nucleotides are characteristic of RDR2 strands

*In vivo*, P4R2 RNAs and siRNAs frequently have 3’ nucleotides that are mismatched to the DNA template and of unclear origin (Blevins et al., 2015; Zhai et al., 2015). Analysis of M13 transcripts revealed that mismatched nucleotides are characteristic of RDR2 transcript 3’ ends, regardless of transcript length (Figure 6A). For Pol IV transcripts, 3’ mismatched nucleotides occur at the same frequency as 5’ mismatched nucleotides and represent background levels typical of RNA-seq reads.

**Figure 6.**
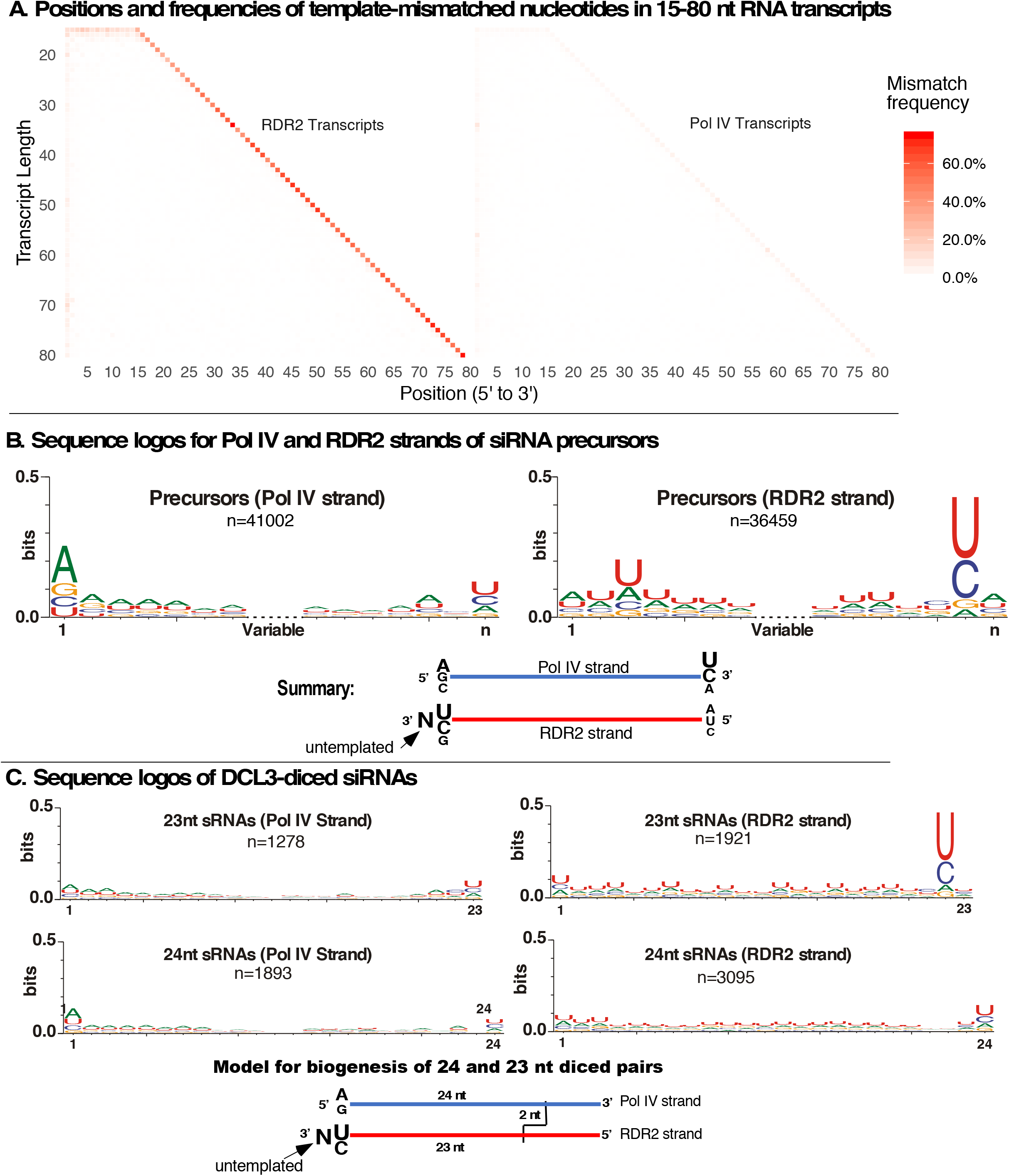
RDR2 transcripts have untemplated 3’ terminal nucleotides. **A.** Positions and frequencies of template-mismatched nucleotides within Pol IV and RDR2 transcripts generated from M13 DNA. Transcripts of 15-80 nt were examined. The frequency of mismatches to the corresponding DNA sequence at each position is denoted in shades of orange/red. **B.** Consensus sequences for unique Pol IV and RDR2 transcripts longer than 26 nt. The 7 nt nearest the 5’- and 3’ ends are shown. Redundant sequences were condensed into unique sequences prior to analysis. Sequence logos were generated using WebLogo 3.6. In the summary cartoon, N denotes the untemplated nucleotide ascribed to RDR2 terminal transferase activity. **C.** Sequence logos for Pol IV and RDR2 strands of 23 and 24 nt DCL3-diced siRNAs. Summary cartoons at the bottom propose models to account for 23 nt RDR2 strand-derived siRNAs with 3’ terminal untemplated nucleotides (left) or 24 nt RDR2 strands whose 3’ ends are complementary to Pol IV transcript 5’ ends.

Among RNAs longer than 26 nt, Pol IV transcripts generated from M13 DNA tend to initiate with a purine, in agreement with prior studies (Blevins et al., 2015; Zhai et al., 2015), with A>G (Figure 6B). Pol IV strand 3’ ends show some enrichment for U>C (Figure 6B). RDR2 strands have a weak preference for A at their 5’ ends, consistent with the weak preference for U at the 3’ end of complementary Pol IV strands (Figure 6B). However, the most striking feature of RDR2 strands is the strong U/C consensus one nucleotide upstream from the 3’ end (Figure 6B). We deduce that this strong U/C signature is complementary to the strong A/G signature of Pol IV strand 5’ ends, with the extra, DNA-mismatched nucleotide present at the extreme 3’ ends of RDR2 strands resulting from an untemplated nucleotide added after dsRNA synthesis is complete. We further deduce that RDR2’s terminal nucleotidyl transferase activity (Blevins et al., 2015) is responsible for addition of the untemplated nucleotide. Sequence data indicate that addition of more than one untemplated nucleotide is rare (Figure S5).

Examination of diced Pol IV-RDR2 precursors generated in the M13 system shows that untemplated 3’-terminal nucleotides are greatly enriched among 23 nt siRNAs derived from the RDR2 strand (Figure 6C), adjacent to the strong U/C consensus at the penultimate position. This suggests a model whereby DCL3 cuts the Pol IV strand 24 nt from its 5’ end, creating a 2 nt 3’ overhang with respect to the RDR2 strand. If the paired RDR2 strand was extended 1 nt at the 3’ end by RDR2’s terminal transferase activity, the resulting RDR2 strand siRNA is 23 nt (Figure 6C, bottom left). Duplexes can also be cut from the other end, generating 24 nt siRNAs that correspond to the 5’ end of RDR2 strands (see Figure 5C). The alternative ways in which precursor duplexes can be diced presumably dilutes the A/G consensus signature of Pol IV transcript 5’ ends, making this signature less prominent among diced siRNAs.

## Discussion

Pol IV and RDR2 represent a unique partnership between DNA- and RNA-dependent RNA polymerases whose concerted reactions convert the information of single DNA strands into double-stranded precursors of 24 nt siRNAs. *In vivo*, it has been unclear which enzyme synthesizes which strand of dsRNA precursors (Blevins et al., 2015; Zhai et al., 2015). Our results show definitively that Pol IV synthesizes first-strand RNAs and that Pol IV’s unusual sequence-independent mode of termination, induced by template-nontemplate strand basepairing, is needed to channel Pol IV transcripts to RDR2 for second strand synthesis.

Multisubunit RNA polymerases generally terminate in response to RNA signals (Porrua et al., 2016). In *E. coli*, RNA stem-loops followed by a string of uracils, or RNA-encoded Rho protein recruitment sites mediate most termination events (Ray-Soni et al., 2016). In eukaryotes, sequence-specific cleavage of nascent transcripts enables RNA binding by termination factors that induce Pol I and Pol II termination (Birse et al., 1997; Connelly and Manley, 1988; El Hage et al., 2008; Goodfellow and Zomerdijk, 2013; Kuehner et al., 2011). Pol III termination is induced by strings of uridines in nascent transcripts, reminiscent of rho-independent termination in *E. coli* (Arimbasseri and Maraia, 2015; Nielsen et al., 2013). Interestingly, the thymidines of the nontemplate DNA strand, complementary to the uridines of the nascent transcript, have been shown to play a role in Pol III termination (Arimbasseri and Maraia, 2015). In light of these other case studies, the Pol IV termination mechanism appears to be distinct, being dependent on the extent of basepairing between the DNA template strand and nontemplate DNA, in cis or in trans, and being sequence-independent. A possibility is that Pol IV simply cannot displace more than 12-18 nt of nontemplate strand DNA, possibly due to weak motor activity associated with the lack of a conserved trigger loop (Landick, 2009).

An intriguing hypothesis had suggested that Pol IV misincorporation errors might induce Pol IV termination, especially at methylcytosines (Zhai et al., 2015), thus accounting for template-mismatched nucleotides at the 3’ ends of P4R2 RNAs *in vivo* (Blevins et al., 2015; Zhai et al., 2015). Pol IV transcription is error-prone (Marasco et al., 2017), however our current study shows that untemplated nucleotides are found almost entirely at the 3’ ends of RDR2 strands, not Pol IV strands. Moreover, methylated cytosines in CG, CHG, or CHH contexts, in the template strand, the nontemplate strand, or both, have no discernable effect on Pol IV termination (Figure 4 D) or Pol IV misincorporation frequency *in vitro* (Marasco et al., 2017). These results argue against the Pol IV-misincorporation hypothesis and point to RDR2’s terminal transferase activity as the likely explanation for the 3’ untemplated nucleotides of P4R2 RNAs and siRNAs.

Our results also argue against the hypothesis that Pol IV transcripts are short because of the high methylcytosine density of Pol IV-transcribed regions, proposed to cause misincorporation and Pol IV termination, if uncorrected. Instead, our data are consistent with a simple model (Figure 7) in which Pol IV initiates within the context of a ~20-30 nt transcription bubble, like other multisubunit RNA polymerases (Bae et al., 2015; Barnes et al., 2015; Holstege et al., 1997; Kahl et al., 2000). Pol IV then transcribes one strand to the edge of the bubble, encounters basepaired DNA and can extend only 12-18 nt more before terminating, accounting for short transcripts that average ~32 nt *in vivo*.

**Figure 7.**
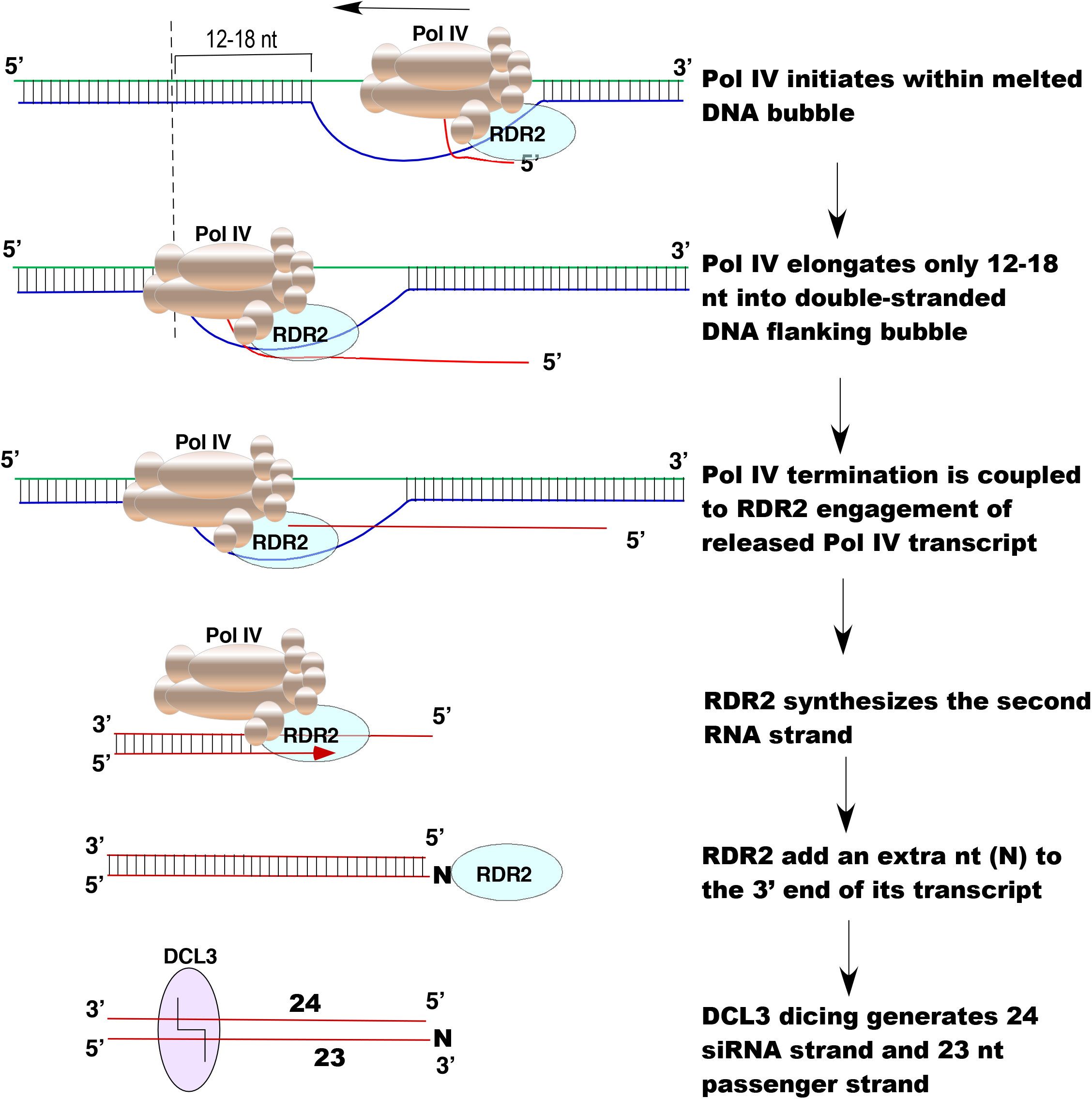
Model for the sequential reactions of Pol IV, RDR2 and DCL3 that account for siRNA biogenesis and RNA channeling in the RNA-directed DNA methylation pathway.

Pol IV termination induced by encountering nontemplate DNA is critical for RDR2 to engage the free 3’ end of the released Pol IV transcript and use it as a template, synthesize the complementary strand and then add an additional untemplated nucleotide to the 3’ end (see Figure 7). This untemplated terminal nucleotide is enriched among 23 nt siRNAs in the M13 system (Figure 6), as in vivo (Wang et al., 2016) and is present at the same end of precursor duplexes as Pol IV strand 5’ ends (see Figure 7). Pol IV strands labeled at their 5’ ends give rise to labeled 24 nt siRNAs following dicing (Figures 5B,C). This suggests a model in which DCL3 measure 24 nt from the 5’ end of the Pol IV strand and cuts to leave a 2 nt 3’ overhang, like other Dicers (Park et al., 2011). Due to the extra untemplated nucleotide at the 3’ end of the RDR2 strand, the resulting DCL3-cleaved RDR2 strand is 23 nt. Importantly, siRNAs found in association with AGO4 are almost exclusively 24 nt (Havecker et al., 2010; Qi et al., 2006). This leads us to propose that 23 nt RNAs typically serve as the passenger strands for 24 siRNAs loaded into AGO4. The terminal transferase activity of RDR2 may thus contribute to passenger strand/siRNA strand discrimination (Figure 7), with the enzymatic properties of Pol IV, RDR2 and DCL3 accounting for RNA channeling from initial transcription to Argonaute loading.

## Acknowledgments

JS dedicates this work in loving memory of his father, S. Tejinder Pal Singh. The authors thank Jeremy Haag for valuable contributions at the onset of the work, Michele Marasco for pioneering the use of M13 for dsRNA synthesis, and the Center for Genomics and Bioinformatics and the Drosophila Genome Resource Center at Indiana University for RNA sequencing and facilities for insect cell culture, respectively. This research was supported by NIH grant GM077590 and funds to CSP as an Investigator of the Howard Hughes Medical Institute. JS was supported, in part, by a Carlos O. Miller fellowship (Indiana University). HYH was supported by NIH NRSA award F32GM125334.

## Author Contributions

VM generated recombinant RDR2 and DCL3 and performed the experiment of Fig. 1D. HYH generated RNA-seq libraries. FW performed bioinformatic analyses of Fig. 5E and Fig. 6. JS performed all other experiments. CSP and JS wrote the manuscript.

## Materials and Methods

### Plant genotypes

*Arabidopsis* thaliana plants used in the study were all of the Col-0 genetic background. Transgenic line genotypes used for Pol IV and/or RDR2 affinity purification were: NRPD1-FLAG *nrpd1-3*, NRPD1-FLAG *nrpd1-3 rdr2-1* and RDR2-HA *rdr2-1 nrpd1-3*. Leaves of 3-4 week old plants were flash frozen in liquid nitrogen and stored at −80° C.

### Affinity purification of proteins

Four grams of frozen leaf tissue was ground in liquid-nitrogen using a mortar and pestle, resuspended in 14 mL of extraction buffer (20 mM Tris-HCl pH-7.6, 150 mM sodium sulfate, 5 mM magnesium sulfate, 20 μM zinc sulfate, 1 mM PMSF, 5 mM DTT, and 1X plant protease inhibitor cocktail (Sigma), passed through two layers of Miracloth and centrifuged at 18,000 x g for 15 min to pellet debris. The supernatant was incubated with 25 μL of anti-FLAG M2 or anti-HA resin (Sigma) for 2.5 hours. The resin was pelleted by centrifugation at 200 x g for 2 min and washed twice with 14 mL of extraction buffer minus protease inhibitors. The resin, with associated affinity-captured proteins, was then suspended in 50 μL of 20 mM HEPES-KOH pH-7.6, 100 mM potassium acetate, 5 mM magnesium sulfate, 10% v/v glycerol, 20 μM zinc sulfate, 0.1 mM PMSF, 1 mM DTT.

### Synthetic nucleic acids used in transcription assays

DNA and RNA oligonucleotides used in the study were purchased from Integrated DNA Technologies, Inc. and are listed in Table 1.

### DNA template-RNA primer hybridization

Equimolar amounts of template DNA and RNA primer oligos were mixed in annealing buffer (30 mM HEPES-KOH pH 7.6, 100 mM potassium acetate), brought to a boil in a water bath and slowly cooled to room temperature. For reactions involving nontemplate DNA, a 10% excess of non-template oligo was included in annealing reactions. RNA primer end-labeling was accomplished using T4 polynucleotide kinase (T4 PNK, NEB) and 25 μCi of ATP, [*γ*-^32^P]-6000 Ci/mmol (Perkin Elmer).

### In-vitro transcription

Transcription reactions used 50 μL of affinity resin slurry with associated Pol IV, RDR2 or Pol IV-RDR2 (or non-specifically associated proteins of non-transgenic plant controls) mixed with transcription reaction buffer to bring the final volume to 100 μL. Final concentrations of reaction components were 20 mM HEPES-KOH pH-7.6, 100 mM potassium acetate, 60 mM ammonium sulfate, 10 mM magnesium sulfate, 10% v/v glycerol, 20 μM zinc sulfate, 0.1mM PMSF, 1mM DTT, 0.8U/μL Ribolock™ (Thermo Fisher). Reactions involving end-labelled primer RNA included 25nM each of the DNA template and primer and 1 mM each of ATP, GTP, CTP and UTP. For body labelling of RDR2 strands, 250 nM T-less template DNA and 1 mM each of GTP, CTP and UTP, 40 μM ATP, 10 μCi of ATP, [α-^32^P]-3000 Ci/mmol (Perkin Elmer) was used. Transcription reactions were incubated 1h at room temperature on a rotating mixer, then stopped by addition of 25 mM EDTA and incubation at 75° C for 10 min. Transcription reactions were then passed though PERFORMA™ spin columns (Edge Bio) according to the manufacturer’s protocol, adjusted to 0.3 M sodium acetate (pH 5.2). 15 μg Glycoblue™ (Thermo Fisher) was added and RNAs precipitated with 3 volumes of isopropanol at −20°C overnight. Following centrifugation, pellets were washed with 70% ethanol, resuspended in 5 μL of 2X RNA loading dye (New England Biolabs) and heated 5 min at 75° C. RNAs were resolved on 15% polyacrylamide 7M Urea gels. Gels were transferred to filter paper, vacuum dried and subjected to autoradiography or phosphorimaging using a Typhoon™ scanner (GE Healthcare).

### Transcript release assay

*In-vitro* transcription assays were carried out as described above. At various timepoints, transcription was stopped by EDTA addition and resin bearing bound Pol IV-RDR2 complexes was pelleted by centrifugation at 200 x g for 1 min and the supernatant collected. The pellet was washed once and then resuspended in 100 μL of transcription buffer. Following heat treatment at 75°C, supernatant and pellet fractions were precipitated and subjected to PAGE as described above.

### RNase sensitivity assays

RNase-treated transcription reactions involved incubation with 2.5 units of RNase H (New England Biolabs), 2.5 units of RNase I (Promega), or both enzymes, at 37° C for 30 min. For RNase III tests 1 unit of enzyme (Epicentre) was added 10 min after addition of RNases I and H. Reactions were stopped by adding SDS to a final concentration of 0.15% (w/v). For RNase H-only tests, reactions were stopped with EDTA and incubated at 75° C for 20 min.

### Capping reactions

Transcripts were alcohol precipitated and pellets washed twice with 70% ethanol. RNA was resuspended and incubated 1 hr, 37°C with vaccinia virus capping enzyme (New England Biolabs), as per the manufacturer’s protocol, in the presence of 100 μM unlabeled GTP and 10 μCi of [α-^32^P]-GTP, 3000 Ci/mmol (Perkin Elmer).

### Cloning, expression, and purification of recombinant DCL3

A DCL3 cDNA, codon optimized for protein expression in insect cells and including an N-terminal 10X His-tag and C-terminal FLAG and Strep tags, was synthesized by GenScript^®^ and cloned into pUC57. The cDNA was subcloned into a pFastBac™ HT B vector (Thermo Fisher Scientific) and used to make recombinant bacmid DNA in *E coli* DH10Bac cells, according to the supplier’s protocol (Bac-to-Bac^®^ Baculovirus expression system; Thermo Fisher Scientific). DCL3-expressing virus was obtained by celfectin-mediated transfection of bacmid DNA into the sf9 cells, using a multiplicity of infection (MOI) of 2. Cells were grown for 72 hr at 27 °C, collected by centrifugation at 350 x g,10 min, and lysed in hypertonic lysis buffer (50 mM HEPES-KOH pH 7.5, 400 mM NaCl, 10% glycerol, 1mM PMSF, 1% protease inhibitor cocktail). The lysate was centrifuged at 39,200 x g at 4°C, 30 min. The supernatant was incubated with ANTI-FLAG^®^ M2 affinity beads (Sigma-Aldrich) on a rotating mixer for 2 hrs at 4°C. Beads were collected by centrifugation for 5 min at 200 x g, and washed 3X with 20 volumes of lysis buffer and 1X with 20 volumes of elution buffer: 50 mM HEPES-KOH pH 7.5, 150mM NaCl and 10% glycerol, 250 μg/mL 3x FLAG peptide (APExBIO). The eluted fraction was concentrated using a centrifugal filter unit (EMD Millipore) with a 30KDa cutoff size and analyzed by electrophoresis on a 4-20% gradient SDS-PAGE gel and immunoblotting, using anti-DCL3 and anti-FLAG^®^ M2 (Sigma Aldrich) antibodies. Recombinant DCL3 was stored at −20°C in storage buffer (HEPES-KOH pH7.5, 150mM NaCl and 50% glycerol).

### RDR2 activity assays

Recombinant RDR2 transcription was carried out in 100 μl reactions containing ~200 ng RDR2, 25 nM end-labeled 37 nt template RNA, 25 mM HEPES-KOH pH 7.5, 2 mM MgCl2, 0.1 mM EDTA, 0.1% Triton X 100, 20 mM Ammonium acetate, 3% PEG 8000, 0.1 mM of rATP, rGTP, rCTP and rUTP, respectively and 0.8 U/μL RiboLock (Thermo Fisher Scientific). Reactions were incubated at room temperature, 60 min. Biotinylated template RNA was incubated with 250 ng streptavidin or 50 μl streptavidin-agarose resin (Thermo Fisher Scientific). Reactions were stopped with 10 mM EDTA, and RNAs purified by TRIzol extraction and ethanol precipitation. Reaction products were resolved using 15% native PAGE gel electrophoresis and visualized by autoradiography.

### DCL3 dicing

Post-transcription dicing reactions containing 100 ng DCL3, 50 mM HEPES-KOH (pH 7.6), 150 mM sodium chloride, 5 mM magnesium chloride, 5 mM ATP, 0.5 mM GTP and 10% glycerol, in 50 μL reactions, were incubated 60 min at room temperature followed by EDTA addition and incubation at 75° C, 10 min. To detect diced body-labeled Pol IV strands, transcription involved the T-less template and inclusion of 10 μCi α-^32^P-UTP (3000 Ci/mmol) UTP (Perkin Elmer).

### Deep Sequencing and analysis of M13-templated RNAs

RNA transcript and DCL3 cleavage product libraries prepared using TruSeq Small RNA Library Prep Kits (Illumina) were subjected to 80 cycles of paired-end sequencing using a NextSeq 500 instrument. Raw data were processed using 3’ adapter trimming script ‘PE_trimadapter.py’. In brief, if 5’ ends of forward and reverse reads are complementary, and remaining 3’ ends are adapter sequences, reads were merged and treated as single-end sequences. Merged sequences <15 nt were discarded. Remaining paired reads lacking adapter sequences were kept in paired format. For Pol IV and RDR2 transcripts, processed reads were first aligned to the *A. thaliana* TAIR10 genome using Bowtie version 1.2.2 (Langmead et al., 2009), allowing 0 mismatches, to remove any contaminating RNAs. For recombinant DCL3 digestion products, reads mapping to a *Spodoptera frugiperda* draft genome assembly (WGS Project: NJHR01) were similarly removed. Bowtie options *-a-v 0* were used for single-end sequences and options *-a -v 0 --allow-contain* used for paired-end reads. Filtered reads were then aligned to M13mp18 (Bayou Labs) allowing up to 3 mismatches, with option *-a -v 3* for single-end sequences and *-a -v 0 --allow-contain* for paired-end reads. Alignment outputs from single-end sequences and paired-end reads were then merged into one file. Sequences present in both diced and un-diced Pol IV-RDR2 transcription samples were removed to enrich for DCL3 cleavage products.

RNAs analyzed in supplemental figures were single-end sequenced for73 cycles, and 3’ adapter trimming performed using Cutadapt version 1.9.1 (Martin, 2011), with options -a TGGAATTC --discard-untrimmed -e 0 -m 15 -0 8. Processed reads were then analyzed as described above.

The strandedness of aligned sequences was determined by the Flag values in the SAM output. The ‘MD:Z’ field from the alignment output was used to determine positions and numbers of mismatches between reads and the reference. Fractions of reads containing mismatched nucleotides at all positions were calculated. To prepare sequence logos for Pol IV and RDR2 transcripts, the corresponding sequences of the 5’-most and 3’-most 7 nt were prepared by WebLogo 3.6 (Crooks et al., 2004) with options *--format eps --size large --color-scheme classic --errorbars NO*. The sequence logos representing 5’ and 3’ sequence features were then merged into one logo. Sequence logos for 23 nt and 24 nt DCL3 cleavage products were prepared similarly.

The RNA-seq data discussed in this publication have been deposited in NCBI’s Gene Expression Omnibus and are accessible through GEO Series accession number GSE126086 (https://www.ncbi.nlm.nih.gov/geo/query/acc.cgi?acc=GSE126086).

