## Supplemental Information for "Reaction mechanisms of Pol IV, RDR2 and DCL3 drive RNA channeling in the siRNA-directed DNA methylation pathway"

| Table S1. DNA and RNA oligonucleotides used in the study | | | |
|---|---|---|---|
| **A.** | **DNA Oligonucleotides** | **Sequence** | **Related to figure/s** |
| 1) | Standard Template DNA | CTAATTCGAGTCAGTCAACGAAAGCTGACTGTGTACGC CTGGTCCGACTCG | 1C, E, F,2A, B, C, 3A, 4A, D, 5B |
| 2) | Non-Temp DNA (16nt) | TACTGACTCGAATTAG | 2A |
| 3) | Non-Temp DNA (25nt) | TCTTTCGTTGACTGACTCGAATTAG | 2C |
| 4) | Non-Temp DNA (28nt) | GCAGCTTTCGTTGACTGACTCGAATTAG | 2A, B, C, 3A, 4A, D, 5B |
| 5) | Non-Temp DNA (31nt) | GAGTCAGCTTTCGTTGACTGACTCGAATTAG | 2C |
| 6) | Non-Temp DNA (36nt) | CACACAGTCAGCTTTCGTTGACTGACTCGAATTAG | 2A |
| 7) | T-Less Temp DNA | CAAAAACGAGACAGACAACGAAAGCAGACAGAGAACG CCAGGACCGACACG | 3B, 4B, 4C, 5C, S2 |
| 8) | T-Less Non-Temp DNA (27nt bp + 5' 1nt unpaired) | GCTGCTTTCGTTGTCTGTCTCGTTTTTG | 3B, 4B, 4C, 5C, S2 |
| 9) | T-Less Non-Temp DNA (27nt bp + 5' 6nt unpaired) | AAAAAGCTGCTTTCGTTGTCTGTCTCGTTTTTG | 4B |
| 10) | T-Less Non-Temp DNA (27nt bp + 5' 11nt unpaired) | AAAAAAAAAAGCTGCTTTCGTTGTCTGTCTCGTTTTTG | 4B |
| 11) | T-Less Non-Temp DNA (27nt bp + 5' 16nt unpaired) | AAAAAAAAAAAAAAAGCTGCTTTCGTTGTCTGTCTCGTT TTTG | 4B |
| 12) | T-Less Non-Temp DNA (27nt bp + 5' 21nt unpaired) | AAAAAAAAAAAAAAAAAAAAGCTGCTTTCGTTGTCTGT CTCGTTTTTG | 4B |
| 13) | T-Less Non-Temp DNA (27nt bp + 5' 26nt unpaired) | AAAAAAAAAAAAAAAAAAAAAAAAAGCTGCTTTCGTTG TCTGTCTCGTTTTTG | 4B |
| 14) | T-Less Non-Temp DNA (22nt bp + 3' 5nt unpaired) | GCTGCTTTCGTTGTCTGTCTCGTAAAAA | 4C |
| 15) | T-Less Non-Temp DNA (17nt bp + 3' 10nt unpaired) | GCTGCTTTCGTTGTCTGTAAAAAAAAAA | 4C |
| 16) | T-Less Non-Temp DNA (12nt bp + 3' 15nt unpaired) | GCTGCTTTCGTTGAAAAAAAAAAAAAAA | 4C |
| 17) | Template DNA (CHG Methylation) | CTAATTCGAGTCAGTCAACGAAAG/iMe-dC/TGACTGTGTACGCCTGGTCCGACTCG | 4D |
| 18) | Template DNA (CHH Methylation) | CTAATTCGAGTCAGT/iMe-dC/AACGAAAGCTGACTGTGTACGCCTGGTCCGACTCG | 4D |
| 19) | Template DNA (CG Methylation) | CTAATTCGAGTCAGTCAA/iMe-dC/GAAAGCTGACTGTGTACGCCTGGTCCGACTCG | 4D |
| 20) | Template DNA (CG and CHG Methylation) | CTAATTCGAGTCAGTCAA/iMe-dC/GAAAG/iMe-dC/TGACTGTGTACGCCTGGTCCGACTCG | 4D |
| 21) | Template DNA (CG, CHG and CHH Methylation) | CTAATTCGAGTCAGT/iMe-dC/AA/iMe-dC/GAAAG/iMe-dC/TGACTGTGTACGCCTGGTCCGACTCG | 4D |

| | | | |
|---|---|---|---|
| 22) | Non-Temp DNA (CHG Methylation) | G/iMe-dC/AGCTTTCGTTGACTGACTCGAATTAG | 4D |
| 23) | Non-Temp DNA (CHH Methylation) | GCAG/iMe-dC/TTTCGTTGACTGACTCGAATTAG | 4D |
| 24) | Non-Temp DNA (CG Methylation) | GCAGCTTT/iMe-dC/GTTGACTGACTCGAATTAG | 4D |
| 25) | Non-Temp DNA (CG and CHG Methylation) | G/iMe-dC/AGCTTT/iMe-dC/GTTGACTGACTCGAATTAG | 4D |
| 26) | Non-Temp DNA (CG, CHG and CHH Methylation) | G/iMe-dC/AG/iMe-dC/TTT/iMe-dC/GTTGACTGACTCGAATTAG | 4D |
| 27) | T-Less DNA Hairpin (30 bp) | GTGTCTGCTTTCGTTGTCTGTCTCGTTTTTGTCCTTTCAAAAACGAGACAGACAACGAAAGCAGACAGAGAACGCCAGGACCGACACG | 4E |
| 28) | T-Less DNA Hairpin (27 bp) | GCTGCTTTCGTTGTCTGTCTCGTTTTTGTCCTTTCAAAAACGAGACAGACAACGAAAGCAGACAGAGAACGCCAGGACCGACACG | 4E |
| 29) | T-Less DNA Hairpin (24 bp) | TCTTTCGTTGTCTGTCTCGTTTTTGTCCTTTCAAAAACGAGACAGACAACGAAAGCAGACAGAGAACGCCAGGACCGACACG | 4E |
| 30) | Bubble Non-Template DNA | GGATACTTACAGCCATATCAGTTACGCCTACTCCATTCCATCCCGGGTTCGTCCAAGTCGACTACTGGATCCTAGGCAGG | 4F |
| 31) | Bubble Template DNA | CCTGCCTAGGATCCAGTAGTCGACTTGGACGAACCCGGGATGGAATGGAGTATTCGCCGTGTCCATGGCTGTAAGTATCC | 4F |
| B. **RNA Oligonucleotides** | | | |
| 32) | RNA Primer (Standard Template) | rUrGrCrArUrArArArGrArCrCrArGrGrC | 1C, E, F, 2 A, B, C, 3A, 4A, 4D, 5B |
| 33) | RNA Primer (T-Less Template) | rUrGrCrArUrArArArGrUrCrCrUrGrGrC | 3B, 4B, 4C, 4E, 5C, S2 |
| 34) | RNA Primer (8nt bp + 8nt 5' unpaired) | rUrUrUrUrUrUrUrUrGrArCrCrArGrGrC | 4A |
| 35) | RNA Primer (8nt bp + 12nt 5' unpaired) | rUrUrUrUrUrUrUrUrUrUrUrUrGrArCrCrArGrGrC | 4A |
| 36) | RNA Primer (8nt bp + 16nt 5' unpaired) | rUrUrUrUrUrUrUrUrUrUrUrUrUrUrUrUrGrArCrCrArGrGrC | 4A |
| 37) | RNA Primer (8nt bp + 20nt 5' unpaired) | rUrUrUrUrUrUrUrUrUrUrUrUrUrUrUrUrUrUrUrUrGrArCrCrArGrGrC | 4A |
| 38) | RNA Primer (Bubble Template) | rUrUrUrUrUrUrUrUrGrGrArCrArCrGrG | 4F |
| 39) | First Strand RNA (40 nt) | rUrGrCrArUrArArArGrUrCrCrUrGrGrCrGrUrUrCrUrCrUrGrUrCrUrGrCrUrUrUrCrGrUrUrGrUrCrU | S1 |
| 40) | Second Strand RNA (40 nt) | rArGrArCrArArCrGrArArArGrCrArGrArCrArGrArGrArArCrGrCrCrArGrGrArCrUrUrUrArUrGrCrA | S1 |
| 41) | RDR2 Transcription template | rUrArCrArArGrCrGrArArUrGrArGrUrCrArUrUrCrArUrCrCrUrArArGrUrCrCrArArCrArUrA | 1D |

**Figure S1 (related to Figure 3A). RDR2 transcripts have triphosphate groups.**
Pol IV and RDR2 transcripts generated using the T-less template were tested for sensitivity to Terminator™ exonuclease, which degrades RNAs with 5' monophosphate groups but not RNAs with 5' triphosphate groups. End-labeled primer RNA was used to initiate transcription and label Pol IV transcripts in lanes 1-6. Alternatively, incorporation of $\alpha$-32P-ATP was used to body label RDR2 transcripts (lanes 7-12). Pol IV isolated from a *rdr2* mutant was tested in the reactions of lane 1-3 and 7. All other reactions had Pol IV-RDR2. Reactions of lanes 11 and 12 underwent treatment with RNA pyrophosphohydrolase (RppH, New England Biolabs) which converts 5' triphosphates into monophosphates. Reactions of lanes 3, 6,10-12 were heated at 75°C, 15 min to melt dsRNA duplexes into ssRNA. Reactions of lanes 2, 3, 5, 6, 9, 10, and 12 were subjected to treatment with Terminator exonuclease. The white line indicates editing of the original image to crop out a marker lane.

The results show that Pol IV transcripts (without RDR2) are sensitive to Terminator when single-stranded, consistent with their 5' monophosphate ends. In contrast, RDR2 transcripts are Terminator-insensitive unless first treated with RppH. These results support the results of the capping assay in Figure 3, as both assays indicate that RDR2 transcripts have 5' triphosphate groups.

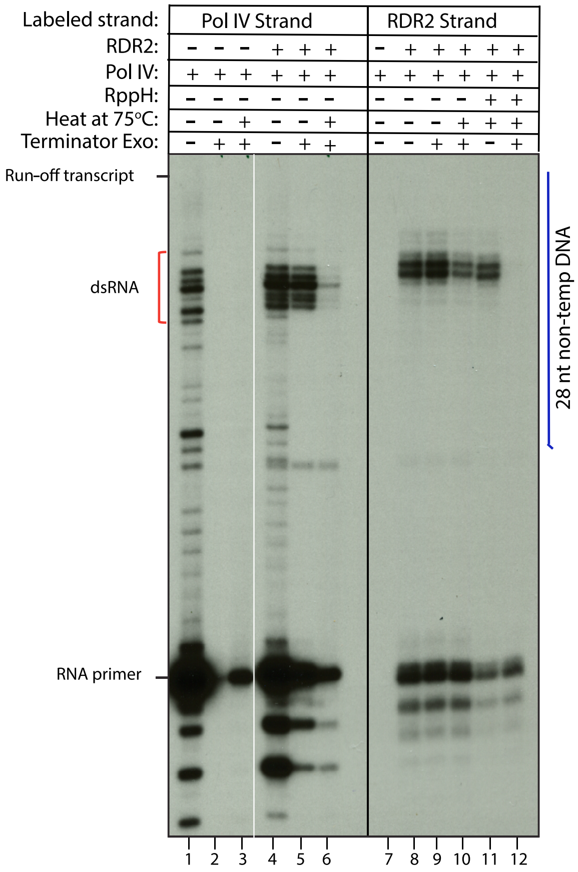

**Figure S2 (related to Figure 3B). Pol IV and RDR2-transcribed strands have different electrophoretic mobilities.**
40 nt synthetic RNAs corresponding to a primer-initiated first strand transcript of the T-less DNA template and its 40 nt reverse complement were end-labeled and subjected to denaturing PAGE and autoradiography. The Table shows the molecular weights of the two oligoribonucleotides. The mass difference is approximately the average mass of a single ribonucleotide.

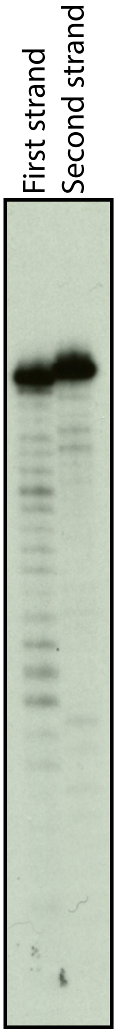


| Oligo | Length (nt) | Mol wt. |
|---|---|---|
| First strand | 40 | 12839.4 |
| Second strand | 40 | 13179 |
| Difference in mol wt. | | 339.6 |

**Figure S3 (related to Figure 5D) Predicted M13MP18 (+) strand folding.** This image, generated using using UNAFold (University of Albany) shows the potential for extensive secondary structure in single-stranded M13 DNA, enabling Pol IV termination and coupling to RDR2 activity.

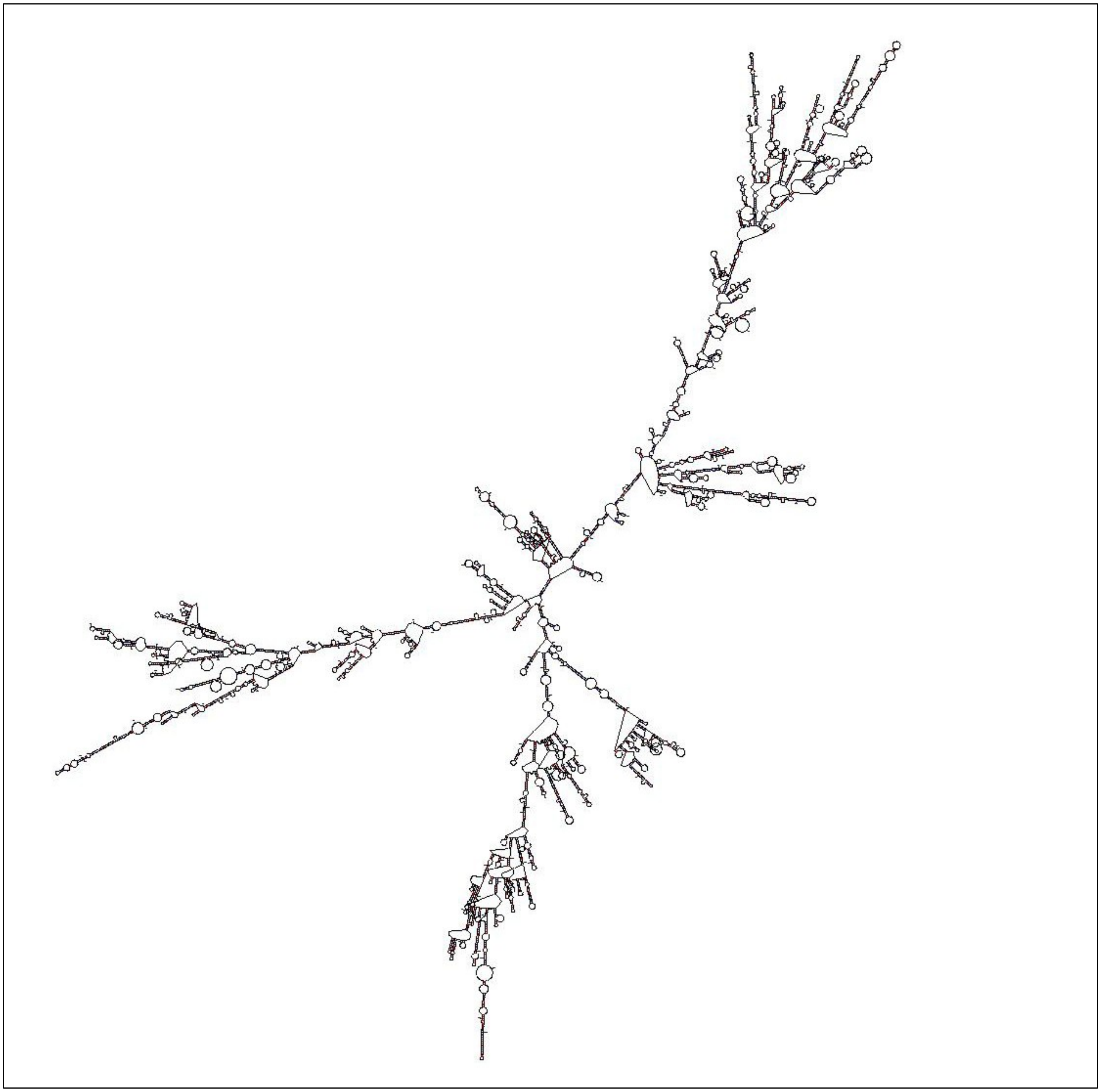

**Figure S4 (related to Figure 5E). RDR2 is needed for second strand transcripts of siRNA precursors generated from M13 DNA**

Second strand transcripts are depleted in the absence of RDR2. The plots show transcript abundance (reads per million) vs. transcript length for first strand (top panel, salmon) or second strand (bottom panel, blue) transcripts, comparing transcripts made by Pol IV-RDR2 complexes (left) to transcripts of Pol IV isolated from *rdr2* mutant plants (right).

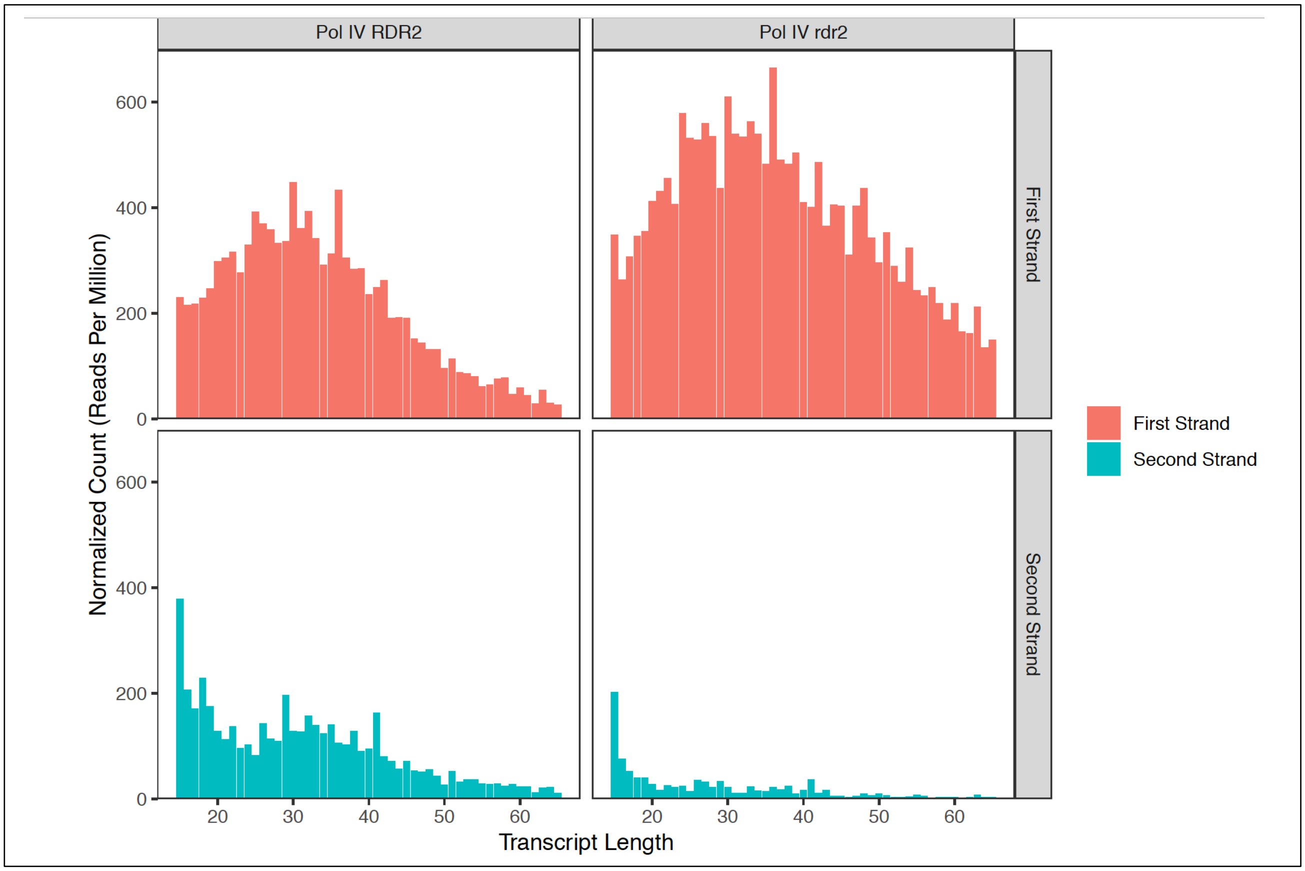

**Figure S5 (related to Figure 6). Features of template-mismatched nucleotides at RNA 3' ends.**

**A.** Number of 3' untemplated nucleotides. The histograms show the frequency of Pol IV or RDR2 transcripts having 0, 1, 2 or 3 template-mismatched nucleotides at their 3' ends.

**B.** Nucleotide identities of 3' untemplated nucleotides. The histograms show the frequency of Pol IV or RDR2 transcripts with single 3' terminal nucleotides (A, U, G or C) mismatched to the DNA template as well as the various sequence combinations for those rare transcripts with two mismatched nucleotides.

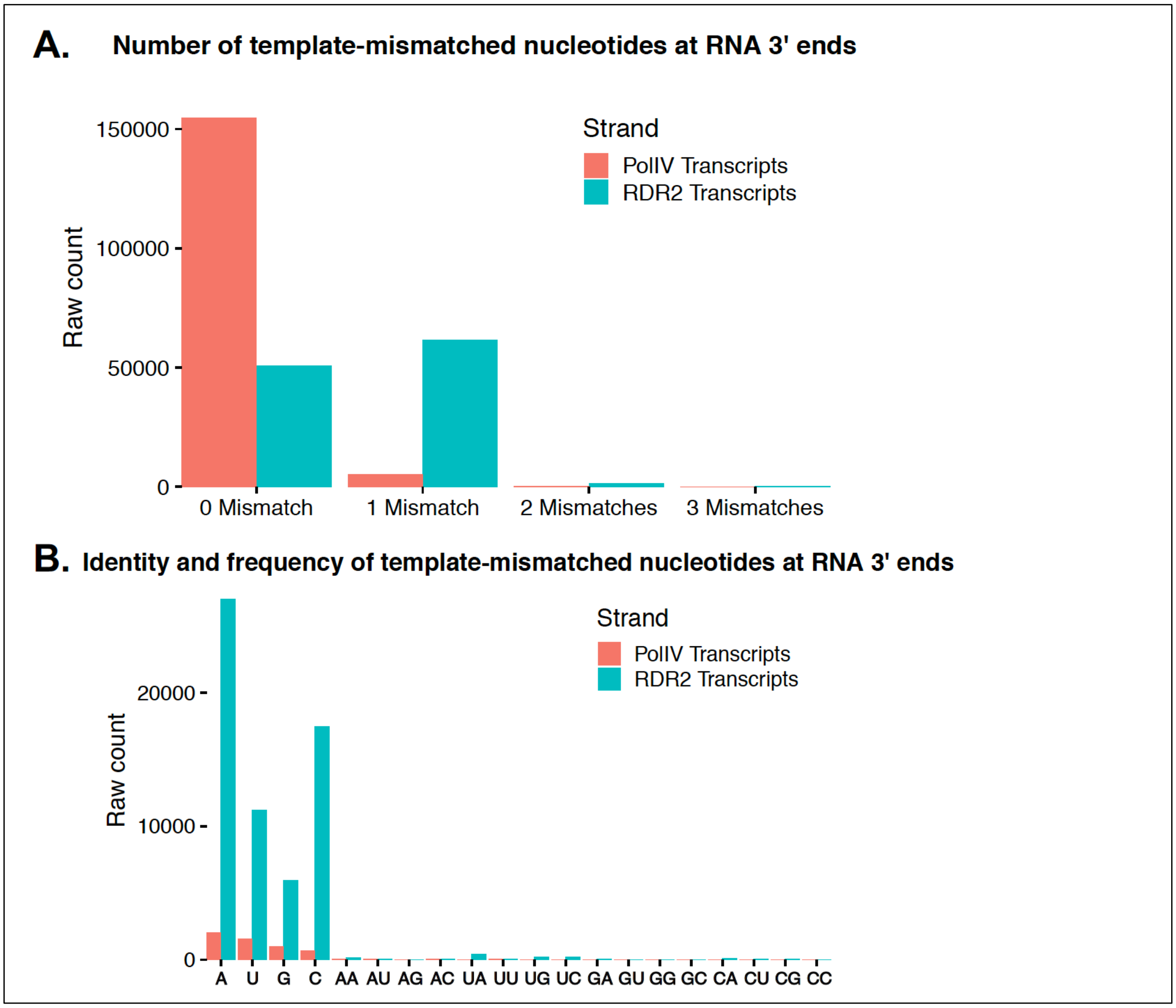
